# Cryptic and not-so-cryptic species in the complex “*Holothuria (Thymiosycia) imaptiens*” (Forsskål, 1775) (Echinodermata: Holothuroidea: Holothuriidae)

**DOI:** 10.1101/014225

**Authors:** François Michonneau

## Abstract

Identifying accurately species is critical for our understanding of patterns of diversity and speciation. However, for many organisms with simple and variable morphological traits, the characters traditionally used by taxonomists to identify species might lead to a considerable under appreciation of their diversity. Recent advances in molecular-data based computational methods have considerably improved our ability to identify and test species limits. Here, we use an integrative approach to delineate species in a complex of sea cucumbers. We used a three-step approach to show that “*Holothuria impatiens*”, a common, shallow-water species, occurring across the Indo-Pacific, the Western Atlantic and the Mediterranean Sea, targeted locally by fisheries, is a complex of at least 13 species. (1) We used the Generalized Mixed Yule Coalescent (GMYC) model to identify putative species without *a priori* hypotheses. In the process, we also show that the number of putative species estimated with GMYC can be affected considerably by the priors used to build the input tree. (2) We assessed based on coloration patterns and distributional information, the most relevant hypothesis. This approach allowed us to identify unambiguously 9 species. However, some of the lineages consistently assigned to belong to different species using GMYC, are occurring in sympatry and are not differentiated morphologically. (3) We used Bayes factors to compare competing models of species assignment using the multispecies coalescent as implemented in *BEAST. This approach allowed us to validate that the species identified using GMYC were likely reproductively isolated. Estimates of the timing of diversification also showed that these species diverged less than 2 Ma, which is the fastest case of closely related species occurring in sympatry for a marine metazoan. Our study demonstrates how clarifying species limits contribute to refining our understanding of speciation.

## 1 Introduction

Despite challenges with definitions, species constitute the biological units used to assess patterns of diversity, identify regions of conservation concern and manage exploited resources [1]. For many taxonomic groups, species limits are misunderstood and the level of undescribed diversity debated, making global estimates of diversity poorly constrained [2, 3]. Additionally, conservation efforts aim at preserving the evolutionary potential of the species, but limited understanding of the processes that maintain present and future diversity hinders these goals [4]. Genetic data, and barcoding in particular, are facilitating the documentation of biodiversity not only by providing an inexpensive and efficient way to identify species and resolve complexes, but also by providing insights into the evolutionary history of the species.

Barcoding has revealed that many species, thought to be well-understood, are complexes. In most cases, these “pseudo-cryptic” species differ morphologically but in traits not traditionally considered, or in traits not available to taxonomists because of the preservation methods or the lack of information about the natural history of the species [5]. For instance, differences in coloration [6, 7], habitat[8], or host preference [9], were not considered as indicative of species limits before cryptic lineages were uncovered by molecular data. In these cases, the single-locus approach provided a way to infer species limits using the evolutionary significant unit (ESU) concept [10] that defines species-level units based on reciprocal monophyly in at least one marker (typically mtDNA), and at least another defining attribute (morphology, distribution, reciprocal monophyly in another trait). If this approach has allowed to unravel high levels of unrecognized diversity, when other defining attributes to infer reproductive isolation are lacking, the patterns of genetic differentiation have remained difficult to interpret objectively.

Many species rely on chemical recognition systems to maintain species integrity (e.g., mate recognition, habitat specificity) [5]. Not only these systems are poorly characterized or difficult to investigate in the context of a taxonomic study, they also rarely translate into morphological, behavioral or ecological differences. The absence of defining traits that correlates with genetic clusters identified from single-locus data make these species “true cryptic” complexes. New methods based on the multispecies coalescent are emerging as powerful ways to investigate species limits in these complexes when evidence is equivocal. By using multi-locus datasets, these methods overcome some of the shortcomings of single-locus analyses that were unable to distinguish between different processes leading to identical patterns (e.g., incomplete lineage sorting and introgression), and do not depend on arbitrary threshold of genetic divergence. Instead, they account for the stochastic process associated with the independent sorting of genealogies along the species tree, and use it to estimate the species tree and the demographic history of the species [11]. These methods also provide a statistical framework to test competing species delineation hypotheses [12].

Among marine organisms, the discovery of high levels of cryptic diversity has challenged the importance of physical barriers as the primary driver of diversification [13]. Since most marine invertebrates have a potentially highly dispersive larval stage, and the oceans in general, and the Indo-Pacific in particular, lack clear geographical barriers to dispersal, opportunities for allopatric speciation seem limited. Yet, while many closely related species have non-overlapping distributions suggesting that allopatric speciation is prevalent, overlap in the distribution of sister species is common. These observations have led to the realization that (1) selection (along environmental gradients, sexual selection, host specificity) may play an important role in reproductive isolation (reviewed in [14]); (2) changes in species distributions have obscured the geographic context at the time of speciation. By clarifying the identity of the species, by providing more accurate estimates of the timing of speciation, and by estimating populations sizes, the multispecies coalescent may shed light on the relative contribution of these factors in shaping patterns of diversity, reveal new patterns and refine our understanding of speciation in the sea.

In this study, we use coloration patterns, genetic, distributional and ecological data, to unravel at least 13 species within “*Holothuria impatiens*” (see column “consensus” in Table 1). This complex includes both species that can be distinguished relatively easily from their live appearance, and species that can only be identified genetically. The broad geographical distribution of this complex and the elucidation of the phylogenetic relationships of its species provide the opportunity to investigate the spatial and temporal dynamics of this radiation.

**Table 1:**
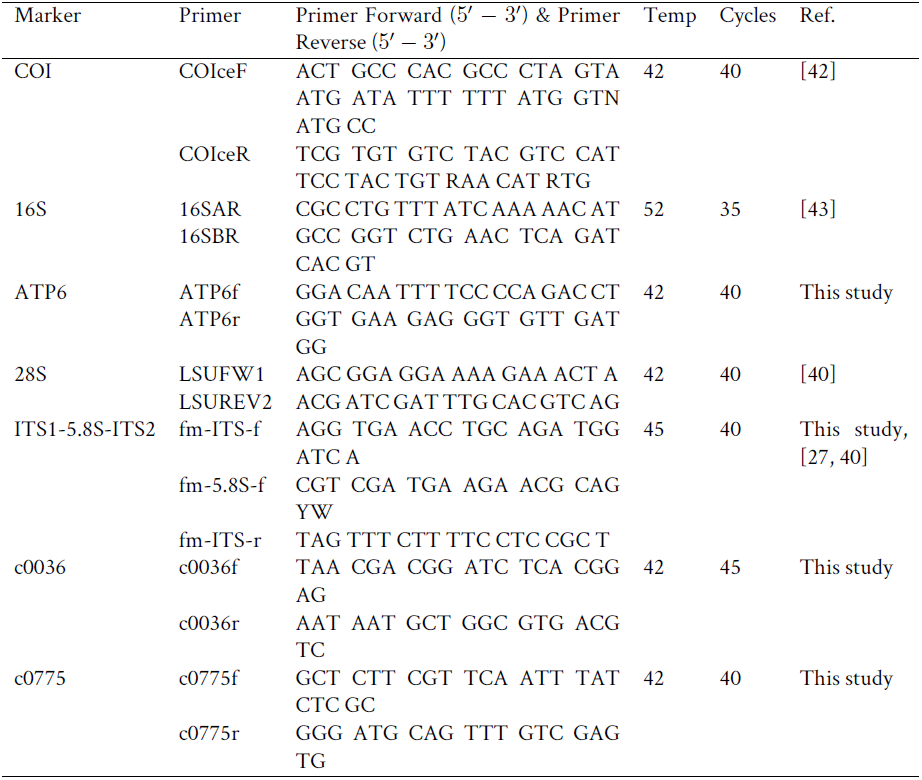
Primers and PCR conditions for the markers used in this study. Temp: annealing temperature, Cycles: number of PCR cycles used to amplify the locus, Ref.: reference used for the primers.

Forsskål described posthumously in 1775 *Fistularia impatiens* from material he collected in Suez, Egypt [15]. The description is limited, but indicates that the body wall is gray with dark spot, and with well-developed, lightly colored tubercles. The drawing accompanying the description corroborates these observations. Since then, the species has been attributed to *Holothuria,* the mostly reef-associated and most diverse genus within the Holothuriidae. The range of *H. impatiens* has been extended, and today is recognized as the most widespread species in the genus. Its range extends from the Red Sea through the entire Indo-West Pacific, to the East Pacific, Caribbean and the Mediterranean. Pearson [16] described *Thymiosycia* as one of five subgenera in *Holothuria*, and designated *Holothuria impatiens* as the type species. The modern concept of *Thymiosycia* was proposed by Rowe [17] to include 13 species.

*Holothuria impatiens* is a common, to locally abundant species found under rocks (in particular in lagoons and back-reef habitats), and is largely restricted to shallow water (< 10 m), although it has been recorded down to 158 m (Y. Samyn, pers. comm.). Because of its ubiquitous distribution, it is one of the more studied species in the family with studies on its reproduction [18]; Cuvierian tubules [19, 20]; toxicity [21]; feeding preferences [22]; parasites [23]; the chemical composition, statistical analysis of the shape, and ontogenic changes in ossicles [24, 25, 26]. It has also been included in molecular phylogenies that investigated relationships among the major groups of holothurians (e.g., [27]) or as an outgroup when studying relationships within a sub-genus (e.g., [28]). *Holothuria impatiens* is a low-value commercial species that is fished in the Eastern Pacific [29], Madagascar [30], and Palau[31].

Despite its relative biological and commercial importance, the variation observed in color patterns reported by previous workers (e.g., [32,p.178], [33]) has yet to be investigated. The goal of this study is to identify cryptic species in the “*Holothuria impatiens*” complex, and to understand the temporal and spatial dynamics of its diversification. To this end, we sampled the entire known geographic range of this complex, and assembled a multi-locus dataset.

To identify species limits in the complex, we used a combination of complementary methods of species delineation following modifications of the approaches outlined by [34, 35, 36]. First, we used the Generalized Mixed Yule Coalescent method (GMYC) on a portion of the mitochondrial locus COI to delineate putative species [37, 38]. We then used independent lines of evidence (color patterns, ecology and geographic information) to assess the validity of these putative species. When no other line of evidence could separate the putative species identified with GMYC, we used model comparison in *BEAST [39, 12] to validate species limits. Our study shows that these methods can be added to the lines of evidence typically used in integrative taxonomy, and provide a powerful tool for evaluating species limits in rapidly evolving groups with limited morphological and genetic differentiation.

## 2 Methods

### 2.1 Sampling

Specimens were collected at low tide, on snorkel, on SCUBA or by dredging. Most specimens were photographed while alive *in situ* or in the lab, anesthetized in a 1:1 solution of sea water and 7.5% solution of magnesium chloride hexahydrate, then preserved in 75% ethanol. When possible tentacles were clipped, immediately put in 95-99% ethanol, and later used for DNA extractions.

Specimens were deposited in the Invertebrate Zoology collections of the Florida Museum of Natural History, University of Florida (UF), Gainesville, FL, USA, while tissue samples are stored in the Genetic Resources Repository of this museum. A few additional tissue samples were taken from vouchers housed at other institutions or were obtained through collaborators without vouchers being retained (Table 5 for details).

**Table 2:**
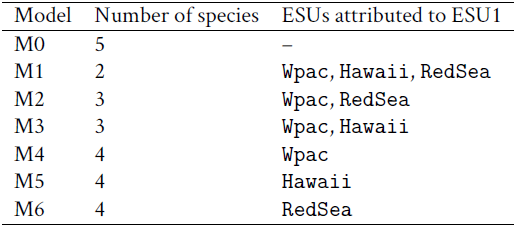
Species groupings used in models of species delineation.

**Table 3:** Characteristics of the loci used for the phylogenetic analyses. *N*: numberof individuals sequenced, *K*: numberof unique sequences, bp: length of the aligned (unaligned) sequences, *S*: number of segregating sites, *Si*. numberof parsimony informative sites. The statistics given for ITS are forthe ones used in the analysis (i.e., after using Gblock).

|  | mtDNA |  |  | nucDNA |  |  |  |  |
| --- | --- | --- | --- | --- | --- | --- | --- | --- |
|  | 16S | COI | ATP6 | c0036 | c0775 | H3a | ITS | LSU |
| $N$ | 94 | 203 | 64 | 70 | 59 | 39 | 61 | 6 |
| $K$ | 87 | 128 | 60 | 57 | 27 | 19 | 58 | 6 |
| bp | 1179 (1117) | 673 (655) | 653 (653) | 637 (624) | 282 (282) | 334 (334) | 1085 (1040) | 1123 (1072) |
| $S$ | 479 | 219 | 330 | 140 | 47 | 46 | 419 | 201 |
| $S_i$ | 383 | 198 | 265 | 97 | 28 | 5 | 237 | 7 |

**Table 4:**
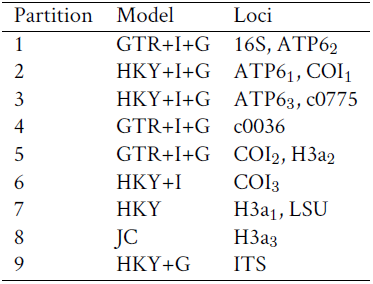
Partition table. The subscript number corresponds to the to the codon position in the alignment. The “real” codon position is offset by 2 for COI, H3a and ATP6 (i.e., COIi is 3rd codon position)

**Table 5:** Specimen information including location, catalog number, and ESU (consensus)

| Catalog Nb. | ESU (consensus) | Location | Specimen Nb. |
| --- | --- | --- | --- |
| MHE006 | EP | Mexico | A0004 |
| MHE048 | EP | Mexico | A0031 |
| FRM134 | EP | Panama | C0028 |
| 3430 | EP | Panama | G0149 |
| FRM106 | EP | Panama | N0132 |
| FRM107 | EP | Panama | N0133 |
| FRM110 | EP | Panama | N0134 |
| FRM124 | EP | Panama | N0167 |
| FRM124small | EP | Panama | N0240 |
| FRM153 | EP | Panama | N0263 |
| 7070a | ESU1 | Mayotte | C0200 |
| 7070b | ESU1 | Mayotte | C0201 |
| 7055b | ESU1 | Mayotte | N1193 |
| 7055b | ESU1 | Mayotte | C0207 |
| 8585 | ESU1 | Lizard Island | FLMNH 036 B03 |
| 8544 | ESU1 | Lizard Island | FLMNH 036 C03 |
| 8459 | ESU1 | Lizard Island | FLMNH 036 E04 |
| 8192 | ESU1 | Lizard Island | FLMNH 036 F07 |
| 8574 | ESU1 | Lizard Island | FLMNH 080 A05 |
| 8604 | ESU1 | Lizard Island | FLMNH 080 C06 |
| 8939 | ESU1 | Tanzania | FLMNH 080 C11 |
| 8603 | ESU1 | Lizard Island | FLMNH 080 G03 |
| 9192 | ESU1 | Europa | FLMNH 160 E05 |
| Samyn | ESU1 | Comoros | K0126 |
| MOLAF125 | ESU1 | GBR | N0023 |
| MOLAF139 | ESU1 | PNG | N0028 |
| 3625 | ESU1 | Palau | N0122 |
| 4343 | ESU1 | Vanuatu | N0123 |
| 2268 | ESU1 | Papua New Guinea | N0125 |
| 3929 | ESU1 | Okinawa | N0127 |
| 3928 | ESU1 | Okinawa | N0128 |
| 4307 | ESU1 | Vanuatu | N0129 |
| 574 | ESU1 | Tikehau | N0130 |
| 4264 | ESU1 | Oman | N0146,C0021 |
| 1850 | ESU1 | Oahu | N0147,C0023 |
| 596 | ESU1 | Moorea | N0148 |
| 10951 | ESU1 | Okinawa | N0553 |
| 10955 | ESU1 | Okinawa | N0554 |
| 8193 | ESU1 | Lizard Island | N0578,FLMNH 036 E07 |
| 8191 | ESU1 | Lizard Island | N0581,FLMNH 036 G07 |
| 10313 | ESU1 | Guam | N0582 |
| 11210 | ESU1 | Marquesas | N1009 |

Table 5: (continued) specimen information
| Catalog Nb. | ESU (consensus) | Location | Specimen Nb. |
| --- | --- | --- | --- |
| 932 | ESU1 | Hawaii | N1149 |
| 5556 | ESU1 | Vanuatu | S0050 |
| 5566 | ESU1 | Vanuatu | S0057 |
| 6485 | ESU1 | Réunion | S0193 |
| 6487 | ESU1 | Réunion | S0195 |
| REU 172 1 | ESU1 | Réunion | S0220 |
| 6774 | ESU1 | Majuro | S0403 |
| 6748 | ESU1 | Majuro | S0404 |
| 6745 | ESU1 | Majuro | S0407 |
| 6957 | ESU1 | Kosrae | S0419 |
| 6989 | ESU1 | Kosrae | S0444 |
| 6990 | ESU1 | Kosrae | S0445 |
| 6988 | ESU1 | Kosrae | S0446 |
| 6987 | ESU1 | Kosrae | S0447 |
| 6982 | ESU1 | Kosrae | S0448 |
| 6967 | ESU1 | Kosrae | S0455 |
| 7081 | ESU1 | Guam | S0464 |
| 4255 | ESU1 | Oman | X0011 |
| 318 | ESU1 | Rangiroa | X0030 |
| 4258 | ESU1 | Oman | X0031 |
| 4098 | ESU1 | Taiwan | X0032 |
| 4257 | ESU1 | Oman | X0033 |
| 5326 | ESU1 | Moorea | X0034 |
| 5294 | ESU1 | Moorea | X0037 |
| 5339 | ESU1 | Moorea | X0039 |
| 5320 | ESU1 | Moorea | X0040 |
| 5206 | ESU1 | Moorea | X0041 |
| 5290 | ESU1 | Moorea | X0042 |
| 5679 | ESU1 | Philippines | X0044 |
| 5597 | ESU1 | Bohol | X0046 |
| 7163a | ESU2 | Nosy Bé | C0202 |
| 7163b | ESU2 | Nosy Bé | C0203 |
| 8406 | ESU2 | Lizard Island | FLMNH 036 B06 |
| 8172 | ESU2 | Lizard Island | FLMNH 036 C07 |
| 8174 | ESU2 | Lizard Island | FLMNH 036 D07 |
| 8355 | ESU2 | Lizard Island | FLMNH 036 G11 |
| 7488 | ESU2 | Nosy Bé | FLMNH 043 A06 |
| 7531 | ESU2 | Nosy Bé | FLMNH 043 B09 |
| 7541 | ESU2 | Nosy Bé | FLMNH 043 D11 |
| 7384 | ESU2 | Nosy Bé | FLMNH 043 G07 |
| 8943 | ESU2 | Tanzania | FLMNH 080 B04 |
| 8944 | ESU2 | Tanzania | FLMNH 080 E05 |

Table 5: (continued) specimen information
| Catalog Nb. | ESU (consensus) | Location | Specimen Nb. |
| --- | --- | --- | --- |
| 9628 | ESU2 | Ningaloo | FLMNH 094 B03 |
| 9611 | ESU2 | Ningaloo | FLMNH 094 B05 |
| BNZ1761 | ESU2 |  | G0348 |
| 216 | ESU2 | Guam | N0124 |
| 220 | ESU2 | Guam | N0126 |
| 10123 | ESU2 | Heron Island | N0485,N1161 |
| 11012 | ESU2 | Okinawa | N0555 |
| 7221 | ESU2 | Nosy Bé | N1155 |
| 7287 | ESU2 | Nosy Bé | N1156 |
| 7318 | ESU2 | Nosy Bé | N1158 |
| 6371 | ESU2 | Réunion | S0214 |
| 7421 | ESU2 | Réunion | S0417 |
| 6983 | ESU2 | Kosrae | S0449 |
| 6711 | ESU2 | Guam | S0458 |
| 6729 | ESU2 | Guam | S0459 |
| 6712 | ESU2 | Guam | S0460 |
| 7076 | ESU2 | Guam | S0466 |
| 6726 | ESU2 | Guam | S0467 |
| 6722 | ESU2 | Guam | S0468 |
| 5271 | ESU2 | Moorea | X0035 |
| 5340 | ESU2 | Moorea | X0036 |
| 5263 | ESU2 | Moorea | X0038 |
| 5625 | ESU2 | Bohol | X0048 |
| 5608 | ESU2 | Bohol | X0049 |
| 7519a | ESU3 | Nosy Bé | C0208 |
| 7519b | ESU3 | Nosy Bé | C0209 |
| 7383 | ESU3 | Nosy Bé | FLMNH 043 A08 |
| 8942 | ESU3 | Tanzania | FLMNH 080 A03 |
| MOLAF138 | ESU3 | PNG | N0027 |
| 5624 | ESU3 | Siquijor Prov. | N0575,X0043 |
| 7360 | ESU3 | Nosy Bé | N0576,FLMNH 043 H05 |
| 8928 | ESU3 | Tanzania | N0577,FLMNH 080 H09 |
| 8941 | ESU3 | Tanzania | N0579,FLMNH 080 G01 |
| 7469 | ESU3 | Nosy Bé | N0580,FLMNH 043 D04 |
| RUMF-ZE-00074 | ESU3 | Okinawa | N0758 |
| 5877 | ESU3 | Yap | S0066 |
| 9637 | Gala | Galapagos Islands | FLMNH 154 A02 |
| 9641 | Gala | Galapagos Islands | FLMNH 154 E02 |
| 9667 | Gala | Galapagos Islands | N0057,N0188,FLMNH 154 G06 |
| 9668 | Gala | Galapagos Islands | N0061,N0185,FLMNH 154 H06 |
| 9655 | Gala | Galapagos Islands | N0184,FLMNH 154 G04 |
| 8336 | gracilis | Lizard Island | FLMNH 036 D01 |

Table 5: (continued) specimen information
| Catalog Nb. | ESU (consensus) | Location | Specimen Nb. |
| --- | --- | --- | --- |
| 8348 | gracilis | Lizard Island | FLMNH 036 D11 |
| 8288 | gracilis | Lizard Island | FLMNH 036 E08 |
| 8354 | gracilis | Lizard Island | FLMNH 036 E11 |
| 8364 | gracilis | Lizard Island | FLMNH 036 F03 |
| 9440 | gracilis | Ningaloo | FLMNH 093 A03 |
| 9544 | gracilis | Ningaloo | FLMNH 093 C06 |
| 9412 | gracilis | Ningaloo | FLMNH 120 B04 |
| 10078 | gracilis | Heron Island | FLMNH 162 E12 |
| 10207 | gracilis | Heron Island | FLMNH 168 B09 |
| 10145 | gracilis | Heron Island | FLMNH 168 C04 |
| 10192 | gracilis | Heron Island | FLMNH 168 D08 |
| 10184 | gracilis | Heron Island | FLMNH 168 F08 |
| 10146 | gracilis | Heron Island | FLMNH 168 G03 |
| 10208 | gracilis | Heron Island | FLMNH 168 G08 |
| 10193 | gracilis | Heron Island | FLMNH 168 H07 |
| 10206 | gracilis | Heron Island | FLMNH 168 H09 |
| 10043 | gracilis | Heron Island | FLMNH 171 A07 |
| 10129 | gracilis | Heron Island | FLMNH 171 C03 |
| 10130 | gracilis | Heron Island | FLMNH 171 D03 |
| 10053 | gracilis | Heron Island | FLMNH 171 D04 |
| 10124 | gracilis | Heron Island | FLMNH 171 G03 |
| 5620 | gracilis | Bohol | N0372 |
| 10122 | gracilis | Heron Island | N0484 |
| 10127 | gracilis | Heron Island | N0556,FLMNH 171 E02 |
| 10126 | gracilis | Heron Island | N0557,FLMNH 171 H01 |
| PH-9 | gracilis | Philippines | N0613 |
| PH-10 | gracilis | Philippines | N0614 |
| PH-11 | gracilis | Philippines | N0615 |
| 8346 | gracilis | Lizard Island | N1159 |
| 4812 | Hawaii | Oahu | C0020 |
| 1858 | Hawaii | Maui | C0022 |
| 1842 | Hawaii | Maui | C0024 |
| 6243 | Hawaii | FFS | FLMNH 023 F07 |
| 6138 | Hawaii | FFS | FLMNH 038 C07 |
| 6140 | Hawaii | FFS | FLMNH 038 D06 |
| 6141 | Hawaii | FFS | FLMNH 038 E06 |
| 6139 | Hawaii | FFS | FLMNH 038 E08 |
| 6142 | Hawaii | FFS | FLMNH 038 F06 |
| 6143 | Hawaii | FFS | FLMNH 038 G06 |
| 4808 | Hawaii | Oahu | G0181 |
| 929 | Hawaii | Hawaii | J0321 |
| 1587 | Hawaii | Hawaii | N0149 |

Table 5: (continued) specimen information
| Catalog Nb. | ESU (consensus) | Location | Specimen Nb. |
| --- | --- | --- | --- |
| 962 | Hawaii | Hawaii | N1150 |
| 6059 | Hawaii | Oahu | N1151 |
| 6279 | Hawaii | Maui | N1152 |
| 6280 | Hawaii | Maui | N1153 |
| 6281 | Hawaii | Maui | N1154 |
| 4805 | Hawaii | Oahu | X0027 |
| 1277 | Hawaii | Maui | X0029 |
| MOLAF087 | Medit | Sicily | G0242 |
| MOLAF123 | Medit | Sicily | N0014 |
| MOLAF124 | Medit | Sicily | X0002 |
| 9699 | RedSea | Egypt | N0583,FLMNH 084 H05 |
| 9701 | RedSea | Egypt | N0584,FLMNH 084 D05 |
| 12030 | RedSea | Djibouti | N1163 |
| 12049 | RedSea | Djibouti | N1164 |
| 12051 | RedSea | Djibouti | N1165 |
| 4713 | tiger | Marianas | FLMNH 112 G06 |
| 4737 | tiger | Guam | G0095 |
| 4709 | tiger | Guam | G0108 |
| 1280 | tiger | Maui | J0419 |
| 5970 | tiger | Maui | S0072 |
| 6929 | tiger | Kosrae | S0441 |
| 11956 | tigerRedSea | Djibouti | N1162 |
| 12086 | tigerRedSea | Djibouti | N1167 |
| 12184 | tigerRedSea | Saudi Arabia | N1168 |
| 7902 | WA | Belize | C0199 |
| 2100 | WA | Virgin Islands | G0222 |
| LACM 2003 76 | WA | Bocas | G0341 |
| 11817 | WA | StMartin | N1006 |
| 11743 | WA | StMartin | N1007 |
| 11790 | WA | StMartin | N1008 |
| 1611b | Wpac | Palau | G0102 |
| PH-HI-1 | Wpac | Philippines | N0627 |
| PH-HI-2 | Wpac | Philippines | N0628 |
| PH-HI-3 | Wpac | Philippines | N0629 |
| PH-HI-4 | Wpac | Philippines | N0654 |
| PH-HI-5 | Wpac | Philippines | N0655 |
| 6771 | Wpac | Majuro | S0402 |
| 6809 | Wpac | Kosrae | S0450 |
| 5610 | Wpac | Bohol | X0050 |

We examined 284 specimens morphologically and 208 were used for molecular analyses. These specimens were collected across the entire known range of *H. impatiens:* Mediterranean Sea, Caribbean Sea, Red Sea, tropical Indian and Pacific Oceans (Table 5).

### 2.2 DNA extraction and amplification

DNA was extracted using either Invitrogen▯ DNAZol® or Omega Bio-Tek^TM^ E.Z.N.A® Mollusc DNA kit following manufacturer recommendations. DNA was most often extracted from tentacles, sometimes from gonads, longitudinal muscles or body wall. When possible, the extractions were performed on tissue sampled in the field.

In this study we amplified the mitochondrial markers COI, 16S, ATP6 and the nuclear markers histone 3 (H3a), 18S, ITS1-5.8S-ITS2, c0036 and c0775 (Table 1). ATP6 primers were developed in this study based on sequences available for this locus in GenBank. c0036 and c0775 are anonymous markers developed from a 454 run on genomic DNA from *Holothuria edulis*, and additional details will be provided in a future study. Briefly, these loci were identified using blastx on the contigs obtained from the 454 run against the predicted proteins for the sea urchin *Strongylocentrotus purpuratus* genome. c0036 matches a portion of the gene encoding for an histone H3-like centromeric protein A-like (locus XP_003723879), with an e-value of 2.10^−43^. c0775 matches a portion of the gene encoding for the protein SFI1 (locus XP_792620), with an e-value of 3.10^−16^.

Primers and PCR conditions used are provided in Table 1. Because of the length of ITS1-5.8S-ITS2, we used the additional sequencing primer fm-5.8S-f. The primer fm-ITS-f is the reverse complement sequence of the primer 18S-1708R (WN-1708R in [27]), and fm-ITS-r is the reverse complement of LSUFW1 (LSU D1,D2 fw1: 56-74 in [40]).

We conducted PCR in 25*μ*L reactions using either 15.4 *μ*L of water, 2.5 *μ*L of Sigma-Alrich ®10X PCR buffer, 2.5 *μ*L of dNTP, 2 *μ*L of MgCl_2_, 1 *μ*L of the forward primer, 1 *μ*L of the reverse primer, and 0.1 *μ*L of Sigma-Aldrich®Jumpstart^TM^ Taq DNA polymerase or the Promega GoTaq MasterMix following manufacturer recommendations.

Sequencing of PCR products was performed by the Interdisciplinary Center for Biotechnology Research at the University of Florida. Chromatrograms were edited using Geneious [41].

Overall, 84 positions (1.14% of the alignment length, and 0.02% of the total number of nucleotide positions in the alignment) contained heterozygous positions. We used the International Union of Pure and Applied Chemistry (IUPAC) nomenclature codes for these positions in the alignment, which are considered as missing data by all the software used in this study.

### 2.3 Definitions

In this paper we use three concepts: “putative species”, “evolutionary significant units (ESUs)”, and “species”:

- “Putative species” describes the lineages identified by the GMYC method and follows the terminology used by the authors of the method [37, 38].
- “Evolutionary significant units (ESUs)” are defined as reciprocally monophyletic lineages in COI that are also characterized by another independent defining attribute (e.g., coloration, morphology, geographical distribution, reciprocal monophyly in another marker). We use this concept as defined by Moritz [10].
- The “species” are the lineages supported by the model of species delineation that best explains the data as assessed by Bayes factors estimated from the multispecies coalescent in *BEAST. They provide an indirect assessment of reproductive isolation and the species are assumed to not exchange genes, forming independent distinct lineages.

### 2.4 Species limits with GMYC

To delineate putative species based on molecular data, we fit the generalized mixed Yule coalescent (GMYC) model ([37, 38, 44]) to the COI sequence data. This approach attempts to detect transitions in the branching rate of an ultrametric tree corresponding to the expected increase in lineage accumulation resulting from the shift between speciation and population-level coalescent events. To determine the location of the threshold, the GMYC model is fit at different nodes along the tree, and the one that provides the best likelihood for the location of the switch is selected. Lineages found beyond this threshold are considered different species.

To obtain the ultrametric trees required by this method, we used BEAST as it provides a powerful statistical framework to interpret the chronological context of molecular variation [45]. Despite some investigations of the effects of priors on the number of species estimated by GMYC ([38, 44, 46]), we here revisit the impact of specifying a strict or a log-normal relaxed clock, as well as the type of tree prior used (coalescent with constant population size [47], coalescent with exponentially growing population size [48], and Yule [49]). The type of clock used will affect branch lengths, it is however difficult to predict *a priori* how the misspecification of this parameter will influence the results. Monaghan et al [38] indicate that using a coalescent constant tree prior is a more conservative approach compared to using a Yule prior, as GMYC uses the coalescent as the null model to explain the branching pattern. Their results validated their predictions and the GMYC method detected more species on trees estimated with a Yule prior. However, these results might be influenced by the nature of the data, in particular by the amount of variation observed in the sequences, and deserve further testing.

To our knowledge few studies have investigated the effect of retaining identical haplotypes in the analysis (but see [46]). In other studies, identical sequences were removed as they were thought to make the task of estimating the transition problematic [38], as the GMYC method cannot accommodate for the null branch lengths associated with identical sequences [44]. Contrary to other programs, BEAST does not assign zero edge length to terminal branches for identical sequences, and thus allows retaining them in the analysis. There are obvious theoretical advantages for retaining identical sequences: BEAST assumes that the sequences represent a random sample of the population when estimating effective population sizes. Removing identical sequences will lead to spuriously high levels of genetic diversity, in turn leading to overestimating population sizes [45, p.98]. As population sizes directly influence the probability that lineages will coalesce, removing identical sequences will lead to longer branches in the trees, blurring the distinction between interspecific and intraspecific coalescent events. We can therefore predict that removing identical sequences of the analysis will lead to higher number of species detected by GMYC and higher uncertainty, both of which are undesirable for the purpose of the analysis.

We performed the GMYC analyses using only COI sequences as the method was designed to be used on single locus datasets, and we wanted to compare how our conclusions might differ if the entire dataset was considered. COI genealogies were estimated using BEAST 1.8.0 [50] and BEAGLE 2.1 [51]. To determine the optimal partition scheme, we used PartitionFinder [52] on the dataset including all sequences as well as on the dataset limited to distinct haplotypes only. We performed a complete search using each codon position as a partition, and limiting the search to the JC, HKY and GTR models with or without a proportion of invariant sites (+I) and/or gamma-distributed rates of evolution (+G) (12 different models *in toto*). Both datasets (all sequences, only haplotypes) favored to assign each codon position to independent partitions. The partition scheme selected by PartitionFinder led to poor mixing due to over-parametrization in initial test runs. We modified the partition scheme slightly, to use: HKY+I+G, GTR+G and HKY for each codon position respectively for the dataset including all sequences; and HKY+G, HKY+G and HKY for the analyses on unique haplotypes. In all analyses, base frequencies were estimated.

We used a log-normal distribution as the prior for the mean rate of substitution for the log-normal relaxed clock and the strict clock, with a log(mean) of-4 and a log(standard deviation) of 1.

As no fossils are known in *Holothuria*, we used the closure of the isthmus of Panama to calibrate the clock. The isthmus of Panama was completely closed about 3 Ma but most species were probably isolated earlier [53]. We chose a conservative estimate to account for the fact that the sister species lineages found across the isthmus are the lineages found in the Galapagos and the Caribbean (the coastal Eastern Pacific lineage is sister to the one found in the Galapagos, Fig. 7). We constrained the age of the node corresponding to the most recent common ancestor of the individuals found in the Galapagos archipelago, the coast of the Eastern Pacific and in the Caribbean using a log-normal prior with a log(mean) of 1.5, a log(standard deviation) of 0.75 and an offset of 2.5. This distribution translates into a mean age of divergence of 6.98 millions years ago (Ma), (95% confidence interval of [3.53,21.99]).

We set the MCMC chain lengths at 5.10^7^, sampling every 5.10^3^ generations. All analyses were run twice from independent starting points and were checked for convergence. From visual inspection of the values sampled by the chains with Tracer 1.6, we determined that a burnin of 10% was sufficient for all runs. ESS values for all parameters were above 350 in each independent run, and the samples from the posterior from each runs were combined.

The trees sampled from the posterior were summarized with treeannotator using the common ancestor tree method, as this method provides a more accurate estimate of the divergence times from the trees sampled from the posterior compared to the other methods available in treeannotator [54].

These trees were used to fit the single-threshold ([37]) and the multi-threshold [38] GYMC model using the R package splits 1.0-19 [55]. We considered the number of “entities” returned by GMYC as the estimated number of species in our analyses. The confidence interval for the number of species corresponds to the minimum and maximum numbers of species for threshold values found within 2 log-likelihood units of the threshold associated with the highest likelihood.

### 2.5 Species limits with coloration and geography

To determine whether the putative species delineated with GMYC were biologically sound, we investigated color patterns, geographical distribution and ecology.

Coloration information was obtained from the photographs of the live specimens and from field observations. Ecological and geographical coordinates were retrieved from the collecting information associated with the specimens.

### 2.6 Species limits with *BEAST

Some of the putative species delineated by GMYC form well defined but shallow clusters, have geographically defined ranges, but cannot be differentiated morphologically. Their ranges overlap (`Hawaii, Wpac`) or border (`RedSea`) the widespread `ESU1`, bringing into question the reproductive isolation of these lineages. To test whether these clusters are reproductively isolated, we compared competing hypotheses of species delineation with *BEAST v 1.8.0 [39] using Bayes factors (BF) estimated by the stepping-stone (SS) and the path sampling (PS) methods [12].

*BEAST infers the species trees from a multi-locus dataset using the multispecies coalescent in a Bayesian framework to account for intraspecies polymorphism and incomplete lineage sorting. This method assumes reproductive isolation (absence of gene flow) among species and requires the assignment of individuals to species *a priori*.

Bayes factors (BF) provide a powerful framework for model selection by comparing marginal likelihoods. The models do not have to be nested and the complexity of the models is directly accounted for by the marginal likelihoods [12]. Kass & Raftery [56] developed guidelines to interpret BF values such that 2.*ln*(*B*_01_) (where 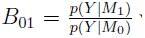 with *y* is the observed data and *M*_*i*_ the model under consideration) must be > 6 to consider the evidence “strong” for (or against) the null hypothesis, and > 10 to consider it “very strong". Here, we used PS and SS methods to estimate and compare the marginal likelihoods *p*(*Y*\*M*) of alternative assignments of individuals to species in *BEAST. With this approach, we tested whether we could detect the signature of reproductive isolation among individuals assigned to different putative species by GMYC but that cannot be teased apart based on color patterns or other evidence.

We investigated whether *BEAST favored models where all or some of the following ESUs were considered distinct species: `WPac, Hawaii and RedSea`, or were panmictic with `ESU1` (Table 2). We included `ESU3` and `gracilis` in our analysis to allow the estimation of the topology when all individuals for the species tested where attributed to `ESU1`. The null hypothesis was that all putative species were reproductively isolated.

In these analyses, we selected individuals for which we had at least one mitochondrial marker and one nuclear marker (43 individuals: 9 of `ESU1`, 7 of `Hawaii`, 7 of `Wpac`, 5 of `RedSea`, 7 of `gracilis`, 8 of ESU3). We used data from the following markers: 16S, COI, ATP6, c0036, c0775, ITS, LSU and H3a; and the same approach as for the phylogeny of the entire complex, using Gblocks to remove ambiguous parts of the alignment in ITS, PartitionFinder to find the most appropriate partition scheme and models of molecular evolution. We unlinked clocks for all markers, used strict molecular clocks, fixing the COI clock at 1%.million years^−1^ which is the rate estimated from the analyses on COI used for the GMYC analyses. The rate for the other clocks were estimated relatively to the COI clock. We ran at least two independent runs for 50.10^6^ generations for each group.

For the estimation of the marginal likelihoods, we used the settings recommended by Baele et al[12] with path steps of 100, and sampling along evenly spaced quantiles of a *Beta*(0.3,1) distribution, but increased the chain lengths to 3.10^6^. We interpreted an absolute difference between the marginal likelihoods estimated with the stepping stone and the path sampling methods higher than 1 as a failure of the estimate to converge. In these cases, we repeated the analysis from a different starting point until the difference between the likelihoods was lower than 1.

To rule out that this approach did not favor models that included more species, we added a “ghost” species in the analysis. This “ghost” species was comprised of individuals that formed a shallow reciprocally monophyletic group in COI within ESU1, but that was not identified as a putative species in any of the sGMYC analyses.

To confirm that COI, which was used to delineate putative species with GMYC, did not drive the differences in marginal likelihoods among the models tested with *BEAST, we repeated some of the analyses omitting COI (models M0 and M4-M6 in Table 2). We also repeated the analyses for these models using only nuclear data to assess the influence of mitochondrial data on the results.

### 2.7 Phylogeny and divergence time estimation

Sequences obtained for each marker were aligned using MUSCLE [57] with the default settings. All markers led to unambiguous alignments except for ITS. For this locus, we used Gblocks [58] to remove parts that were difficult to align using a minimum length of a block of 5 (`-b4=5`), and the option “all gap positions can be selected” (`-b5=a`). We used the resulting alignment to estimate the phylogenetic relationships and the timing of the divergence among the different lineages of *Holothuria impatiens* using RAxML 8.0.1 [59] and BEAST 2.1.2 [60] (with BEAGLE 2.1.2 [51]).

To determine the optimal partition scheme and models of molecular evolution, we analyzed the alignment using PartitionFinder [52]. We defined the data blocks such that the three codon position for ATP6, COI, H3a, and each of 16S, c0036, c0775, ITS and LSU were individual partitions. We used the greedy algorithm, with linked branch lengths, no user tree, and models of molecular evolution available by default in BEAST (JC69, HKY, GTR, without or with a proportion of invariant sites [+I] and/or Gamma distributed rates across sites [+G]). We selected the best-fit models and partition schemes based on the Bayesian Information Criterion. The best fit model had 9 partitions.

For the BEAST analysis, following some initial testing runs, we modified the partition scheme slightly to improve mixing of the chains as the partition scheme selected by PartitionFinder led to over-parametrization (Table 4).

We used three independent strict molecular clocks (one for each of mitochondrial markers, protein-coding nuclear markers, nuclear ribosomal markers) [61]. The tree prior was set to a Yule process, and a random starting tree. The Markov chains were run for 185.10^6^ generations (and sampled every 5.10^3^ generations). The analysis was repeated twice from independent starting points.

We used Tracer 1.6 to check that the MCMC chains had reached stationarity, that mixing was adequate, and that the two independent runs were consistent. We ensured that a representative sample of the posterior distributions was sampled for each independent runs, by checking that the ESS values for all parameters were above 200. The samples from the posterior distribution from both runs were combined using logcombiner after removing a 20% burnin, resulting in 29601 states sampled. The ESS values obtained by combining the runs were all above 440. We summarized the trees sampled with treeannotator using the common ancestor tree method [54].

## 3 Results

### 3.1 Sequence data

The complete sequence data set was 5966 bp (with 2505 bp from mitochondrial genome, and 3461 bp from the nuclear genome, Table 3). Of the 1220 parsimony-informative sites, 69% were from the mitochondrial loci (Table 3).

We obtained sequence data from 207 individuals covering the entire known geographical range of *Holothuria impatiens*. All of these individuals were sequenced for at least one mitochondrial locus, and 36% were sequenced for at least 1 nuclear locus and 34% for 2 or more (Fig. 11).

### 3.2 Species limits with GMYC

The maximum-likelihood estimates for the number of putative species varied widely and ranged from 14 to 20 for the single-threshold method (sGMYC), and from 17 to 23 for the multi-threshold method (mGMYC) (Fig 2).

Species estimates from sGMYC were more conservative and with narrower confidence intervals (CI) than the mGMYC approach. The maximum-likelihood estimates for the sGMYC were in most cases at the lower limit of the CI, while the estimates from the mGMYC were at the upper limit. For the analyses performed on all the sequences, the CI for the estimated number of species using sGMYC and mGMYC did not overlap.

On the genealogies estimated with a strict clock, sGMYC was less sensitive to the type of tree prior used, as the estimated number of species was identical for the three priors tested with 16 and 17 estimated species for analyses on all haplotypes and on unique haplotypes respectively. The Yule prior produced equal or more conservative estimates of the number of species compared to the two types of coalescent priors.

With sGMYC, the estimated number of species was more conservative and less sensitive to the priors used to reconstruct the genealogy when all sequences were included.

Overall, the putative species delineated with sGMYC were consistent across the different priors used. For the analyses based on all sequences, the difference between the 14 species estimated with the Yule prior and the 16 species estimated with the other priors was caused by (1) the lumping of `ESU3 tigerRedSea and tiger;`(2) the later time to coalescence for divergent haplotypes related to `ESU3` with the Yule prior.

For the analyses based on unique haplotypes and a relaxed clock, with the Yule prior, the coalescence of the haplotypes happened later in the tree leading to lower estimated number of species compared to the coalescent priors. With the relaxed clock, many coalescent events were reconstructed in the vicinity of the threshold leading to broad confidence intervals.

With the strict clock analyses, the additional species delineated between the unique haplotypes and all sequences corresponds to the recognition of both `tiger` and `tigerRedSea`.

For all analyses, at least one putative species was represented by a single sequence. A divergent lineage related to `ESU1`, collected in Lizard Island, Australia, was assigned to its own species (`ESU1_Lizard` in Fig. 1). In all analyses (except with the most conservative estimate, i.e., using all sequences, a relaxed clock and a Yule prior), two sequences related to `ESU3` (from the Ryukyus, Japan, and from Papua New Guinea), were each assigned to their own species (`ESU3_Deep` and `ESU3_PNG` respectively in Fig. 1).

**Figure 1:**
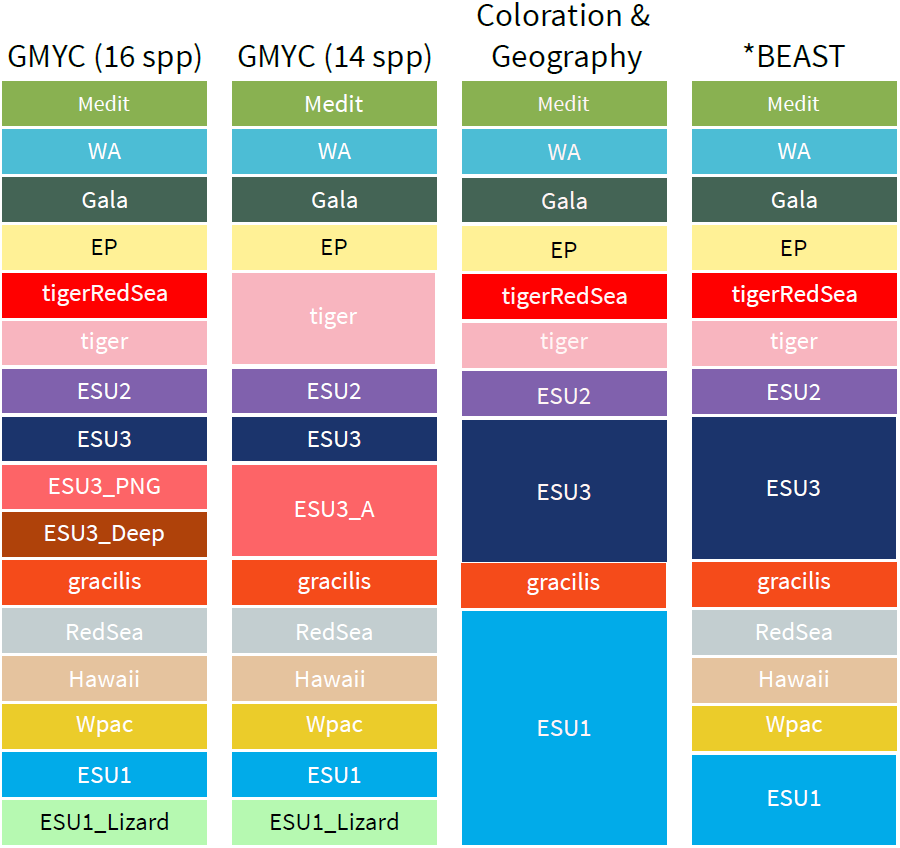
Correspondence between the different criteria used to delineate the species of the “*Holothuria impatiens*” complex. Medit: Mediterranean, WA: Western Atlantic, Gala: Galapagos, EP: Eastern Pacific; PNG: Papua New Guinea; `Wpac`: Western Pacific.

### 3.3 Species limits with coloration and geography

Members of the “*H. impatiens*” complex are characterized by two dominant colors. The dark coloration is typically solid towards the anterior end, forms bands in the middle of the body, and then spots that become rare towards the posterior end. This pattern varies considerably across species and across individuals. The dark color is typically brown but varies in lightness and can take a red or purple tint. The lighter color is typically gray but varies in lightness from light brown to yellow or even white. The light-colored areas typically harbor a “salt-and-pepper” pattern with small dark spots on the light background. Despite these common characteristics, the combination of color patterns and geographical distribution allows us to distinguish unambiguously at least ten species (Fig. 1, 3, 4).

**Figure 2:**
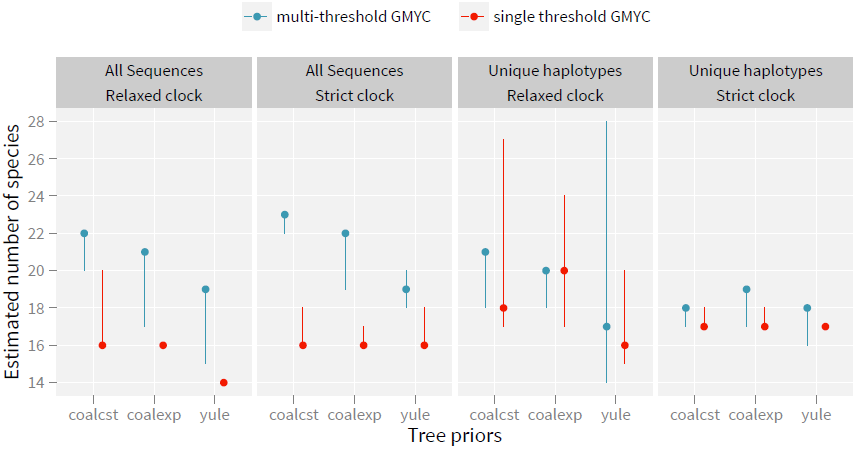
Number of species estimated by single and multi-threshold GMYC methods on COI phylogenies. The dots represent the number of species associated with the node(s) that correspondis) to the highest likelihood(s) for the location of theshift(s), the dotted lines represent the range forthe estimated number of species for nodes within 2 likelihood units of the maximum-likelihood estimate.

**Figure 3:**
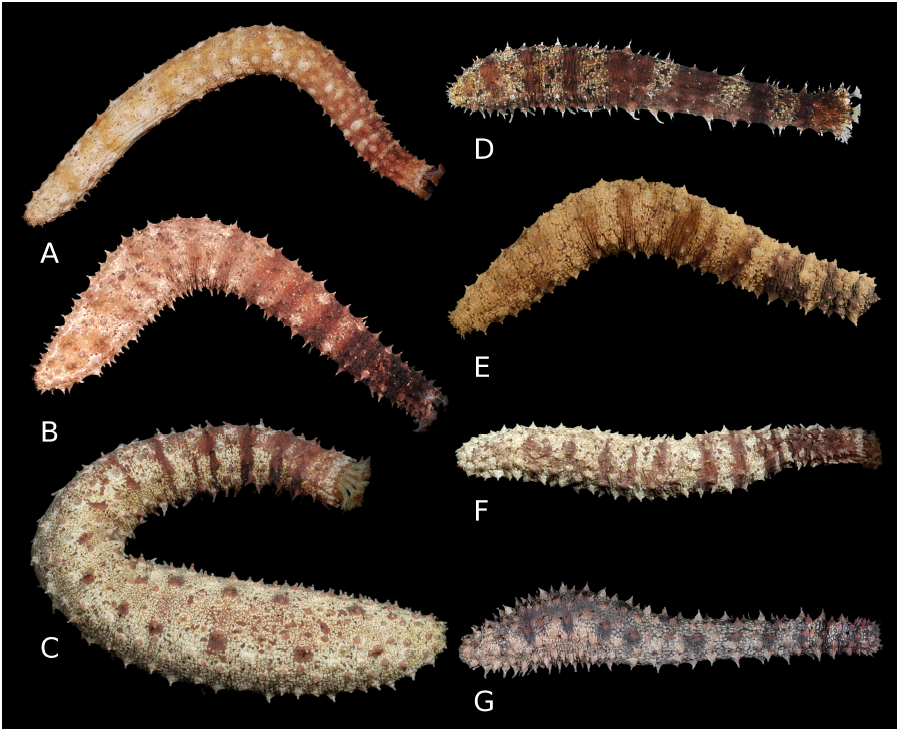
Dorsal view of members of the “*Holothuria impatiens*” complex. A. `ESU1`, Réunion Island, UF6487; B. `ESU1`, Réunion Island, UF6588; C. `ESU1`, Majuro, Marshall Islands, UF6748; D. `ESU1`, Okinawa, Japan, UF10950; E. `RedSea`, Djibouti, UF12070; F. `Wpac`, Majuro Marshall Islands, UF6771; G. `Hawaii`, French Fregate Shoal, `Hawaii`an archipelago, UF6243.

**Figure 4:**
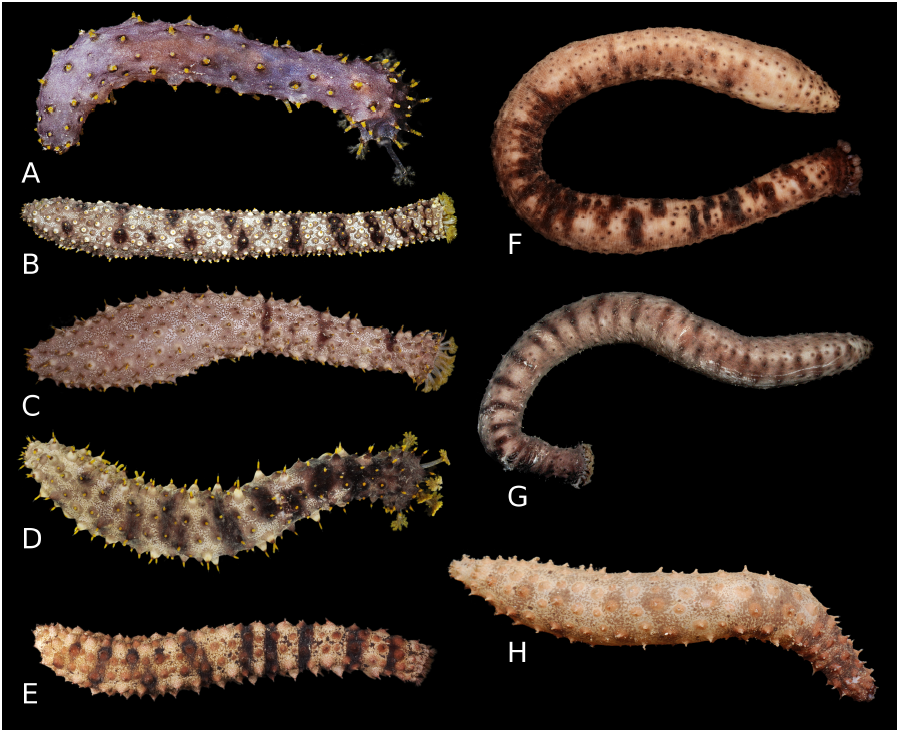
Dorsal view of members of the “*Holothuria impatiens*” complex. A. ESU2, Nosy Bé, Madagascar, UF7287 (juvenile); B. ESU2, Guam, UF6729; C. ESU2, Lizard Island, Australia, UF8355; D. ESU2, Okinawa, Japan, UF11012; E. gracilis, Lizard Island, Australia, UF8288; F. tigerRedSea, Djibouti, UF11956; G. tiger, Kosrae, Federated States of Micronesia, UF6929; H. `ESU3`, Nosy Bé, Madagascar, UF7469.

`Medit, WA, Gala, EP, ESU1` have similar color patterns but subtle differences, and the restriction of their ranges to an oceanic basin or an archipelago allow teasing them apart. `ESU2` is the only species with bright yellow papillae and tentacles (Figure 4). `tiger` and `tigerRedSea` are also very distinctive with sparse, dark, tubercles that barely raise from the body wall, giving them a smoother appearance than the other species. They are also much larger than the other species and specimens reach regularly more than 30 cm. `gracilis` has very large, conical, typically red tubercles, and a very distinctive “gritty” texture. `ESU3` has elongated, pyramidal tubercles with white papillae. However, `ESU1, Hawaii` and `Wpac` have broadly overlapping distributions and are morphologically indistinguishable (Figure 3).

### 3.4 Species limits with *BEAST

Both path sampling (PS) and stepping-stone sampling (SS) gave almost identical estimates of the marginal likelihoods (Fig 6). The values reported for the Bayes factors (BF) are the means of the two independent replicates. Here, the more negative the BF, the stronger the statistical evidence in favor of the null hypothesis specifying that `Wpac, Hawaii, RedSea` and `ESU1` are all reproductively isolated species.

**Figure 5:**
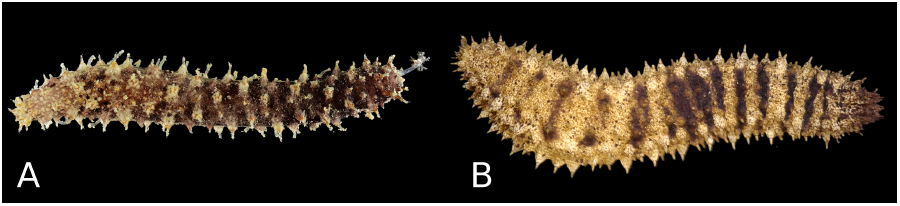
Dorsal view of members of the “*Holothuria impatiens*” complex. A. WA, St Martin, UF11790; B. EP, Panama (no Voucher).

**Figure 6:**
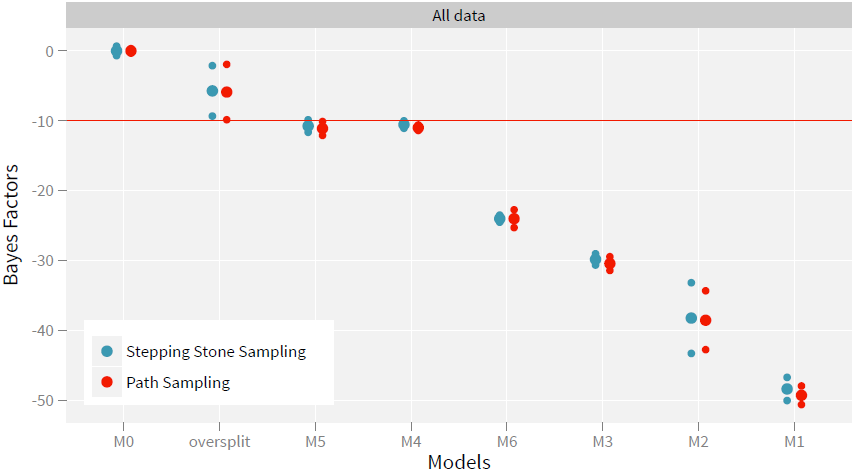
Bayes factors comparing the null hypothesis (M0: ESU1, RedSea, Wpac, Hawaii are all different species) to alternate models of species assignment: additional “ghost” species added (oversplit), no `Hawaii` (M5), no `Wpac` (M4), no `RedSea` (M6), no `Wpac` and no `Hawaii` (M3), no no `Wpac` and no `RedSea` (M2), all individuals attributed to `ESU1` (Ml), see also Table 2. The small dots indicate the values obtained for independent replicated estimations, the large dots the mean of these values. Estimates obtained using the stepping-stone sampling method (on the left in blue) are consistent with the estimates obtained using the path sampling method (on the right in red). The horizontal dashed red line represents the value considered as “very strong evidence” in favor of the null hypothesis according to Kass & Raftery [56].

The model that included putative species identified by GMYC represented by more than one individual, that do not differ morphologically (`Hawaii, Wpac` and `ESU1`) had the highest marginal likelihood (Figure 6). The Bayes factors for the models that considered `Hawaii, Wpac` and `RedSea` as being panmictic with `ESU1` lent “very strong” support in favor of the null hypothesis (BF=–10.8, – 10.5, and –24 respectively). The model with an additional “ghost” species also indicated a poorer fit (BF = –5.7), indicating that this approach did not favor a model that included more species. The model with a random assignment of the species had the worst fit (BF = –243.5).

Similar results were obtained when COI was omitted. The differences in likelihood were however smaller leading to –10 < BF < –6 for models that considered `Hawaii` (BF = –8.2) and `Wpac` (BF = –9.4) as panmictic with `ESU1` indicating “strong” support in favor of the null hypothesis. The BF was still below –10 for the model that considered `RedSea` as panmictic with `ESU1` (BF = –19.9).

Analyses conducted on nuclear data alone led to low BF that do not support the null model (BF = –0.2 for no `Hawaii`, and BF = –4.9 for no `RedSea`). For the model that considered `Wpac` as the same species as `ESU1`, the BF was positive suggesting a better fit to the data than the null hypothesis, but below the threshold to consider the evidence “strong” (BF = 2.8).

### 3.5 Phylogenetic relationships among the ESUs

The phylogenetic relationships recovered with BEAST and RAxML of the species included in the “H. *impatiens*” complex on the concatenated alignment were identical (Fig. 7 and Fig. 12). The topology was highly supported, and the nodes corresponding to the most recent common ancestors of the ESUs had high posterior probabilities (> 0.99) and bootstrap values (10 out of 13 ESUs > 0.9, others > 0.8) for the Bayesian and maximum-likelihood tree respectively.

**Figure 7:**
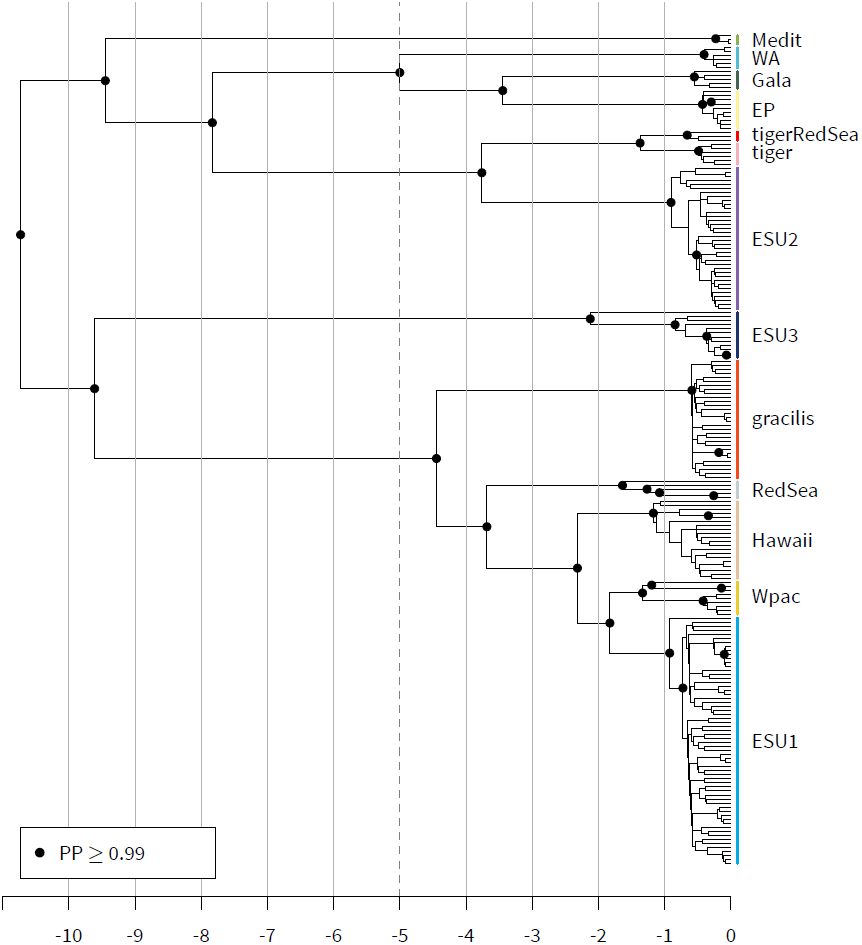
Bayesian phylogenetic tree of the *Holothuria impatiens* complex. Vertical bars along the tips of the tree show the different ESUs found in the complex. Axis on the bottom in million years.

**Figure 8:**
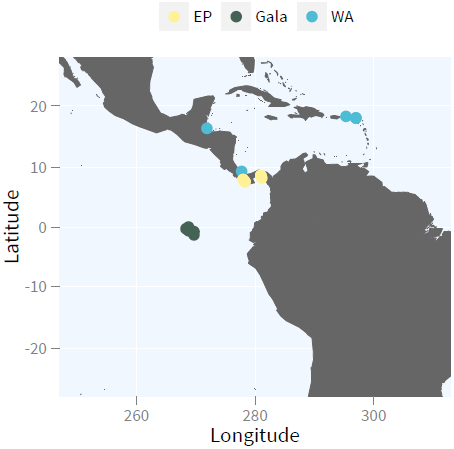
Distribution maps of the ESUs in Eastern Pacific and Caribbean (WA, Gala, EP)

**Figure 9:**
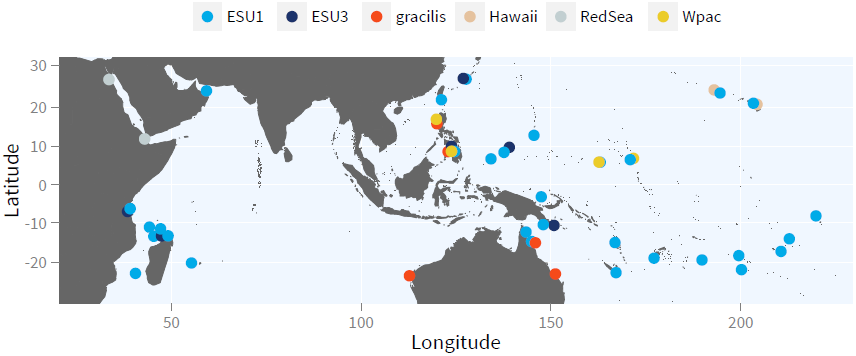
Distribution map for ESU1, ESU3, gracilis, Hawaii, RedSea, Wpac.

**Figure 10:**
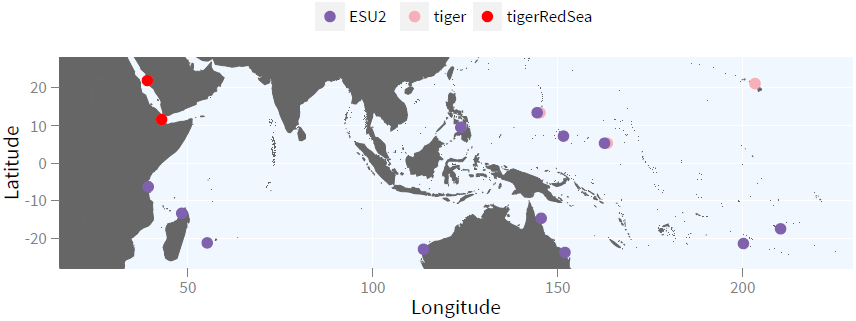
Distribution map for ESU2, tiger, and tigerRedSea.

**Figure 11:**
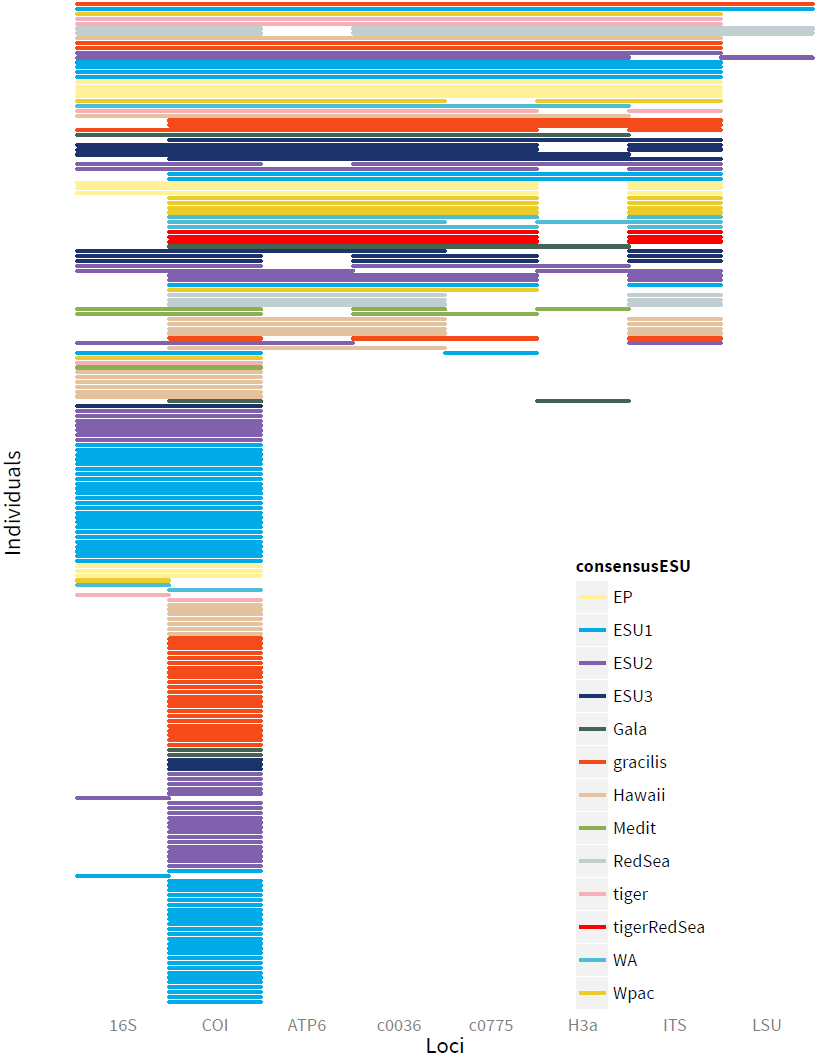
Representation of the loci sequenced for each individual included in the analyses.

**Figure 12:**
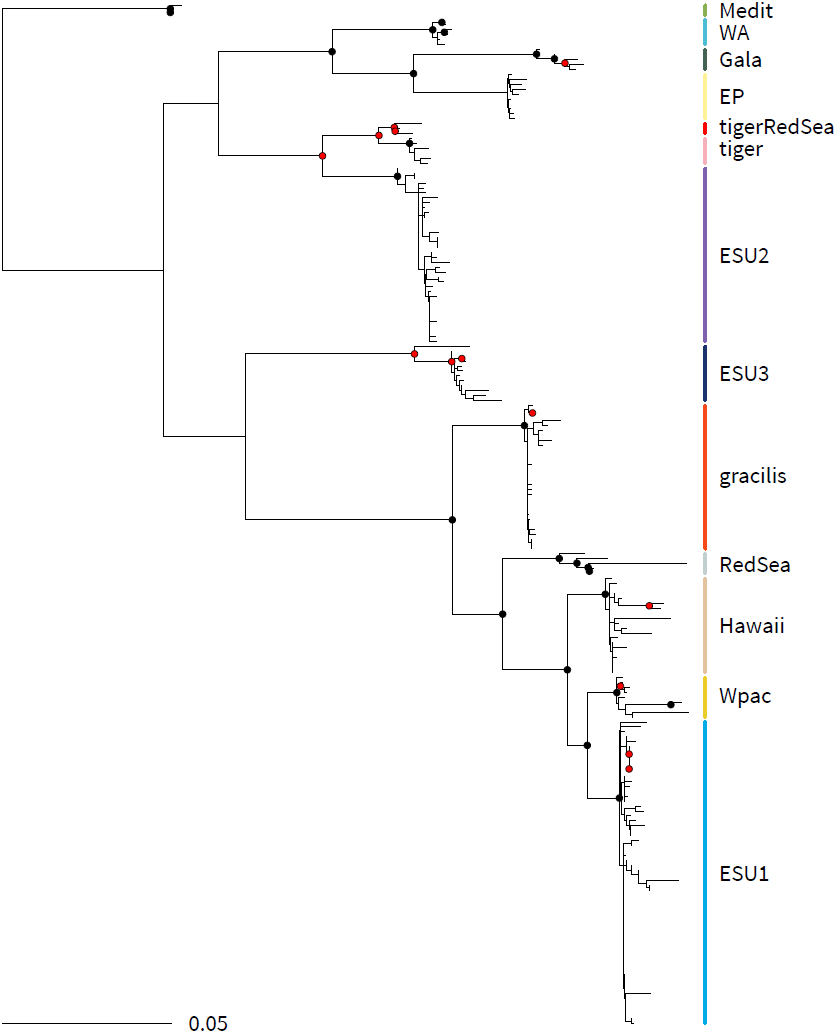
Maximum-likelihood phylogeny for all concatenated loci estimated using RAxML with a GTR+G model for each partition. Black circles represent bootstrap values ≥ 90, red circles bootstrap values ≥ 80. Colored vertical bars represent consensus ESUs.

**Figure 13:**
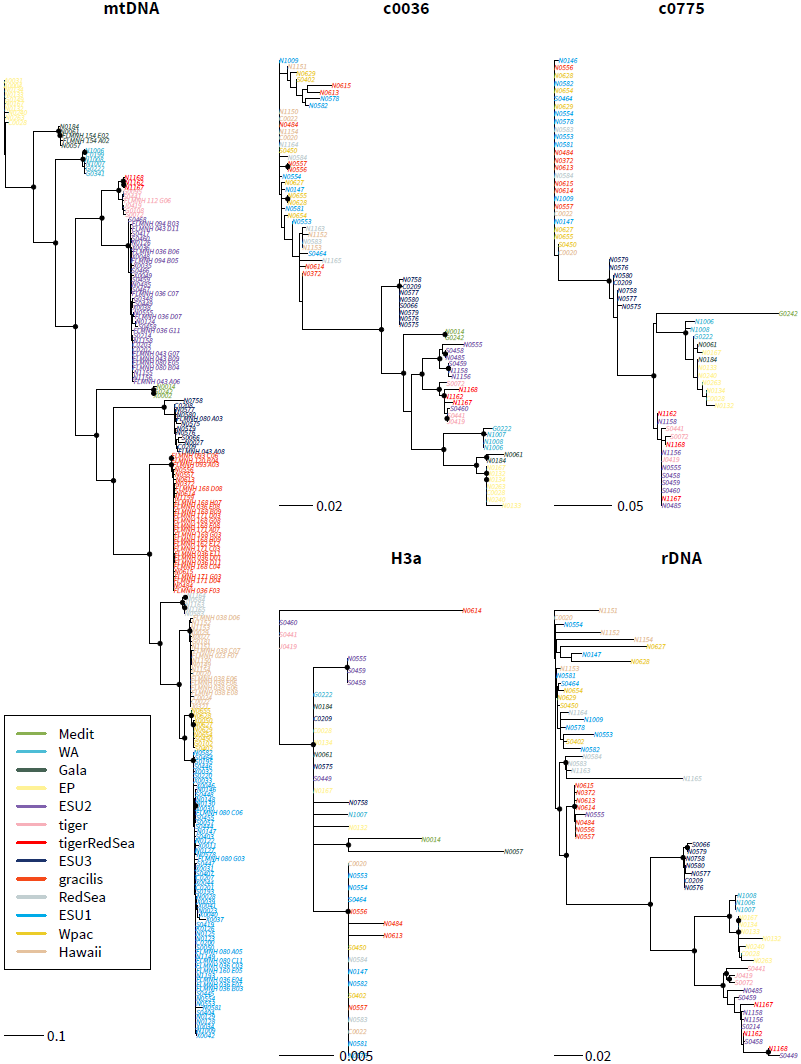
Maximum-likelihood genealogies (unrooted) for each locus reconstructed using RAxML with a GTR+G model of molecular evolution for each partitions. Black circles on the nodes indicate bootstrap values ≥ 80 based on 500 replicates. Terminals are colored based on their ESU. “mtDNA” is the concatenation of ATP6, COI, 16S; “rDNA” is the concatenation of ITS and LSU.

**Figure 14:**
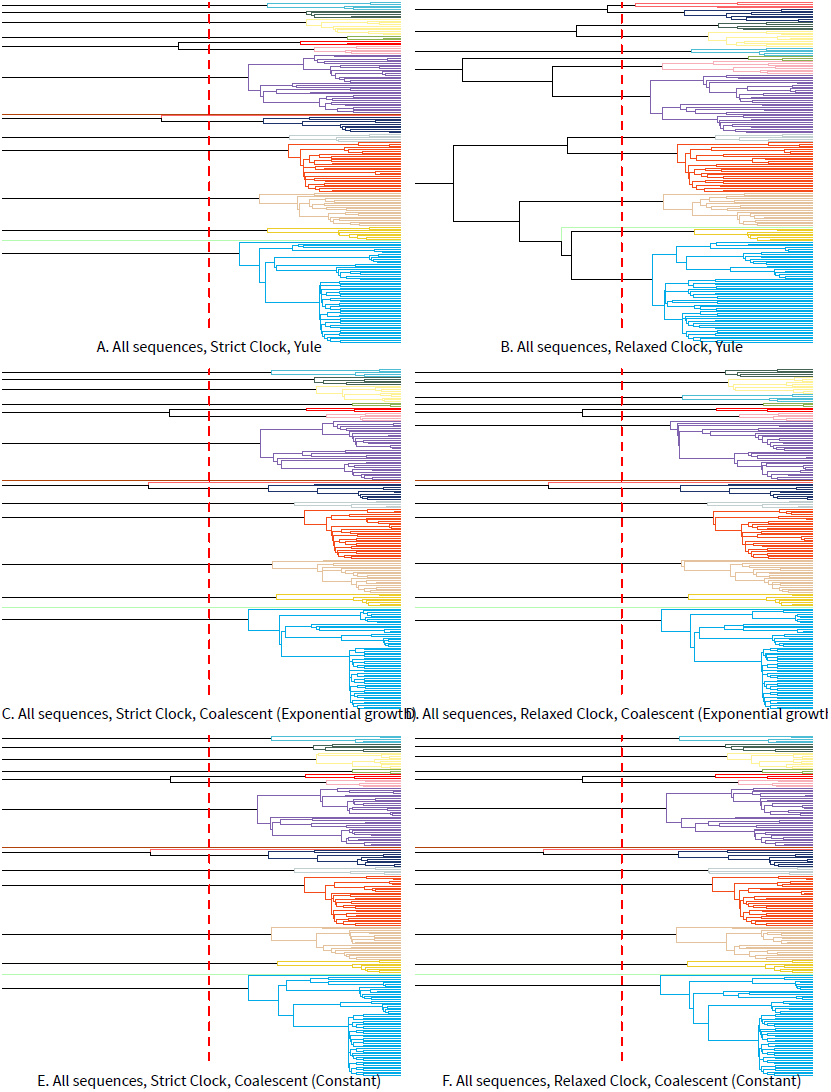
Position of the species delineated with GMYC on COI genealogies estimated using various priors with BEAST. Each color corresponds to a different putative species

**Figure 15:**
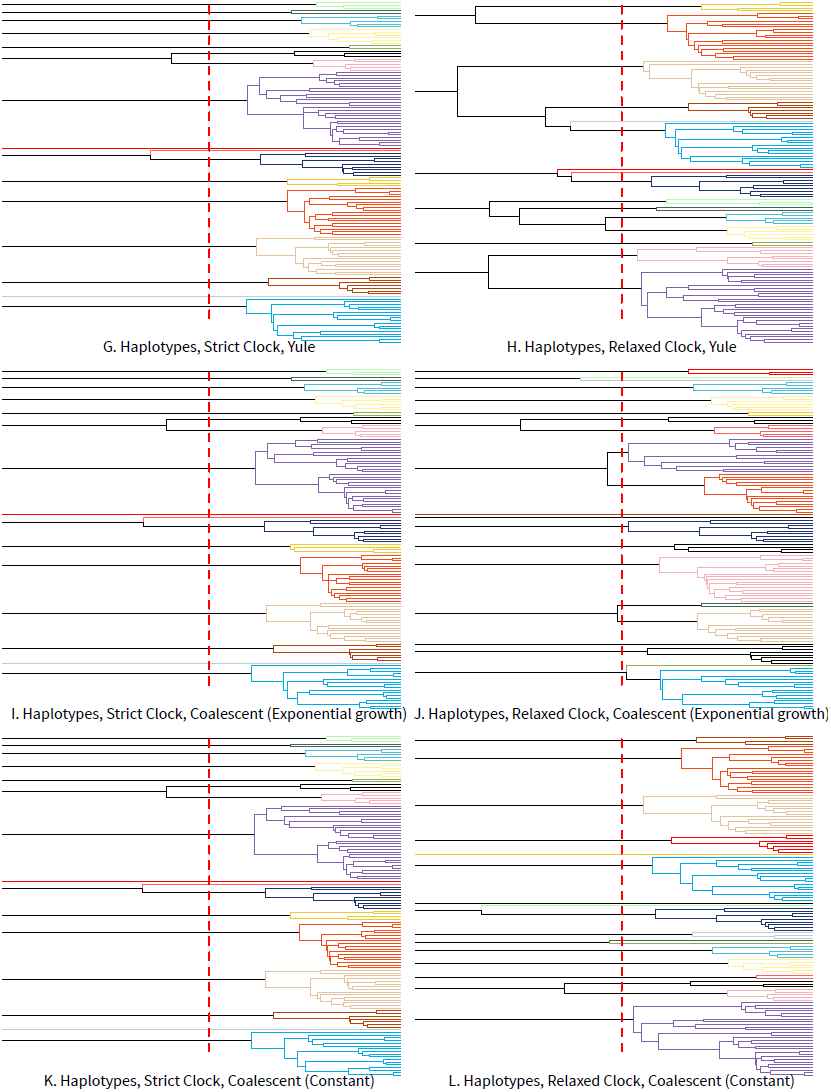
Position of the species delineated with GMYC on COI genealogies estimated using various priors with BEAST. Each color corresponds to a different putative species

Two main clades can be distinguished that diverged approximately 10 Ma (Fig. 7). The first clade (`Medit, WA, EP, Gala, ESU2, tiger, tigerRedSea`) is circumtropical and is found in the Mediterranean Sea, the Caribbean basin, the Eastern-Pacific and across the Indo-West Pacific, including the Red Sea. The other clade (`ESU1, Hawaii, Wpac, ESU3, gracilis`) is restricted to the Indo-West Pacific (including the Red Sea).

The geographical ranges of the species vary considerably. Some are restricted to an ocean basin (`Medit, WA, RedSea tigerRedSea`) or an archipelago (`Hawaii, Gala`) but others have ranges that encompass large sections of the Indo-West Pacific (`ESU1, ESU2, ESU3`). This diversity of distributions results in the co-occurrence of several species of the complex across most of the Indo-West Pacific, and culminates in the Philippines where at least 5 species are found. More generally, at least 3 species are co-occurring in the Northern part of the Western Pacific (the Philippines, Micronesia, Hawaiian archipelago).

Overall, species found in the Indo-Pacific have overlapping ranges, even for species that diverged recently. For instance, `Wpac` and `Hawaii` co-occur with `ESU1` and diverged most recently (1.8 Ma and 2.2 Ma respectively, Fig. 7 and Fig. 9).

## 4 Discussion

### 4.1 Species limits in the “*Holothuria impatiens*” complex

Species delineation is a complex issue and the conclusions will depend of the characters and methods chosen. Here, using complementary approaches, combining discovery (GMYC) and testing methods (*BEAST), we unraveled at least 13 species within the complex “*Holothuria impatiens*” supported by multiple lines of evidence.

The discovery methods, based on genetic differentiation in the mitochondrial locus COI, found putative species that require additional material to be evaluated. In particular, three individuals characterized by divergent haplotypes, related to `ESU1` for one, and to `ESU3` for the others, were consistently recovered as separate species. In both cases, a single individual was available, making it impossible to assess reproductive isolation with *BEAST. The individual related to `ESU1` (UF8191) had the typical color pattern of `ESU1`, with which it co-occurs and no line of evidence seem to indicate that it could be a different species. The same situation occurs with one of the individuals related to `ESU3` (N0027) that cannot be differentiated from other individuals from ESU3. The other individual related to `ESU3` (RUMF-ZE-0074) is small, and was dredged from 32 m in a soft bottom bay of Kume Island, Ryukyus, Japan, a different habitat from the typically reef-asssoicated `ESU3`. The distinct ecology and deeply divergent sequences characterizing this individual suggest it could represent a different species but additional material is needed to assess this hypothesis.

The combination of geography and differences in color patterns, in retrospect, allow the differentiation of 10 species in the complex (Figure 1). Taxonomic studies of most marine invertebrates have relied on preserved specimens. Preservation can render delicate morphological characters, including coloration, difficult or impossible to see, and are frequently lost or fade with time. Because hard parts are easily preserved and their form is generally not impacted by preservation, taxonomists often favor them to differentiate species. For instance, in sea cucumbers, species-level taxonomy has almost entirely relied on the shape of ossicles, microscopic calcareous secretions found in many tissues. Because ossicles show substantial diversity and variability, whilst variation in the simple anatomy of these animals is limited at lower taxonomic levels. Even with ossicles, until recent decades, workers have mostly focused on those found in the body wall, ignoring ossicles in internal structures, and have paid little attention to intraspecific variation, typically illustrating the ossicle assemblage of a single individual. This, in combination with the fact that several of these species co-occur and can share the same habitat, has contributed to the failure to recognize this diversity, and differentiate species in this complex.

Three additional species, that do not seem to be distinguishable morphologically, and that co-occur were validated by the multispecies coalescent statistical framework.

### 4.2 Species limits with GMYC

Despite being widely used to delineate species, the sensitivity of the GMYC method to the factors affecting the shape of the tree used have not been investigated thoroughly. Tree shape can be affected by biological factors or by the type of analysis used to build the tree. Simulated data have been used to test the influence of diversification rates, effective population sizes, and migration rates, on the accuracy and precision of GMYC estimates [62, 63]. Some of the effects of the methodological choices have been evaluated by the authors of the method [37, 38, 44] and by others [46], but the conclusions might be difficult to generalize as they depend in part on the nature of the dataset. Other species delineation methods using sequence data are also available (e.g., RESL [64], ABGD [65], bPTP [66], bGMYC [67]), and it would be interesting to investigate how they perform compared to the results obtained here.

The multiple-threshold method (mGMYC) led to higher estimates of putative species, and was less accurate than the single-threshold method (sGMYC). We did not find any other line of evidence that could corroborate the delineation proposed by this approach. In particular, some of the putative species proposed by mGMYC would split lineages that were otherwise highly supported by the multilocus phylogeny. The overall lower accuracy of mGMYC was also observed by the authors of the method using a simulated dataset. They however found that the 95% confidence set for the estimated number of putative species between sGMYC and mGMYC were overlapping [44]. We observed this pattern on the analyses using only unique haplotypes but not in the analysis that included all sequences.

On our dataset, when analyzing all sequences, the number of putative species was consistently 16, except for the relaxed clock with the Yule prior in which case it was 14. The method is therefore relatively robust to prior specification on our dataset. The 16 species delineation is the most realistic as it recovers morphologically differentiated species that were not considered distinct in the 14 species hypothesis (`tiger` and `tigerRedSea`). Additionally, sGMYC recovered more putative species than we can confidently recognize, since they were represented by single individuals. This is a common pitfall of single-locus species delineation methods, where the broad geographical sampling of many individuals will reveal unique, divergent haplotypes that may not represent different species. However, in our analysis, it is worth noting that all our recognized species were apparently reciprocally monophyletic making the GMYC approach suitable.

The combination of priors used in our study led to overestimating the number of species when it was tested by the authors of the method [38]. On the other hand, Talavera et al [46] did not find any notable differences using trees estimated with different sets of priors. Our dataset differs from these studies by including a younger radiation, less species, and more individuals per species.

Talavera et al [46] did not find that using all haplotypes or only the unique haplotypes affected the number of species estimated with the GMYC method. Here, we confirmed the theoretical expectation that removing the duplicated sequences overestimated the number of species and decreased the precision of the estimates when the tree was estimated with BEAST. Removing identical sequences in the analysis overestimates the effective population size parameters that BEAST uses. In turn, these overestimated sizes spuriously increase the time to coalescence for the lineages, making the estimation of the inflection point between species-and population-level coalescent events earlier in the tree, and more difficult to detect. This bias leads to oversplitting and decreases accuracy. Investigating how many individuals need to be sampled to obtain correct estimates would be worth pursuing.

GMYC was developed with the intent of automating the estimation of the number of species from large-scale single-locus sampling efforts, particularly from environmental samples. In this context, a reasonable amount of inaccuracy or imprecision from the true number of species may be acceptable. The advantage of the method, namely providing a rapid and objective assessment of the number of species, outweighs the possible deviations from the true value. However, in the context of a taxonomic study where the goal is to identify species limits among a few hundred individuals, the factors affecting tree shape need to be considered, and additional lines of evidence need to be included to validate the putative species delineated [68, 63].

### 4.3 Species limits with *BEAST

The statistical framework offered by *BEAST that compares alternative models of species assignment seems a promising venue to investigate species limits, when other lines of evidence are equivocal (e.g., [36, 35]). The requirement to assign specimens to species *a priori* can however be challenging. When morphological evidence is scarce, and deeply divergent lineages are recovered with genetic data, circularity is introduced when the same data is used to recognize species, and to provide statistical support for these species. Here, COI was used to assign species to individuals for putative species that could not be differentiated morphologically.

The confounding effects of using COI to formulate the species hypotheses and to test species limits can stem from: (1) the information contained in the nucleotides forming the locus, and (2) from the absence of recombination in mitochondrial DNA. We attempted to evaluate the relative contribution of these confounding effects by conducting the analyses without COI and without mitochondrial respectively. The results for the analyses conducted without COI were qualitatively similar to the analyses performed with this locus, but the statistical evidence in favor of the hypothesis considering these closely related lineages as different species, was weaker. There was no statistical evidence supporting this hypothesis when the analysis was conducted on nuclear data alone, owing to the low differentiation in these loci for the putative species tested by the models. These results suggest that the remaining of the mitochondrial loci used in our study also favor the null hypothesis, but additional independent nuclear loci with enough differentiation to tease out these species are needed.

Another potential issue with assigning species to individuals based on their COI sequences, is that they may not reflect species limits because of introgression or incomplete lineage sorting. Recently diverged species might still be poly-or paraphyletic in their COI genealogy making species assignment inaccurate. Inaccurate species assignment can potentially be detected from the population sizes estimated by *BEAST that will be unusually high [69, 45]. This would however require to analyze more loci than in the present study to obtain accurate estimates of population sizes [39, 70]. Determining how the Bayes factors will behave if the null hypothesis is misspecified remains to be investigated.

### 4.4 Species limits and coloration

Before the confirmation by molecular methods that differences in coloration were relevant to differentiate closely related species, marine invertebrate taxonomists have been reluctant to consider species defined solely by differences in coloration [5]. Although variation in coloration within “*H. impatiens*” has been noticed by previous workers, and some unusual color morphs were described as sub-species by Clark [32, 71], the focus on ossicle shape to distinguish sea cucumber species has led to an underestimation of the diversity in this group.

The purpose of coloration in sea cucumbers has not been explored but since species can be distinguished based on their color patterns, natural selection may play a role in its evolution. Color patterns have been documented to be diagnostic of species limits in crustaceans [7], nemerteans [5], fish [72]. In some of these cases, the association between color patterns and species limits is maintained by assortative mating (fish [72], crabs [73, 74]). Given that most sea cucumbers inhabit cryptic habitats and do not possess image-forming eyes, the mechanisms maintaining a tight association between coloration and species identity needs to be investigated.

While some species of “H. *impatiens*” harbor slight variations of the typical color pattern characterizing the complex, others deviate from it and are polymorphic. For instance, `ESU2` has two color morphs: a common color pattern that occur throughout the range of the species, and one restricted to the Western Indian Ocean (Nosy Bé, Madagascar and Réunion Island). Both `gracilis` (alternate morph in the southern part of the Great Barrier Reef) and `ESU1` (alternate morph found in Lizard Island, Australia; `Hawaii`; and Guam) are also polymorphic. Interestingly, these alternate color morphs are similar for the three species (uniform chocolate brown background with white to yellowish tubercles), and could result from allelic polymorphism shared across species in genes involved in the pigmentation pathways (as in albinism or melanism), or from convergent selection. These two hypotheses are not mutually exclusive. Furthermore, since coloration seems indicative of species limits, these alternate color morphs could also represent additional cryptic lineages.

Even though, there is no genetic differentiation in the markers used in this study to distinguish individuals exhibiting these alternate color patterns, studies in fish indicate that color differentiation between incipient species might precede any genetic signal (e.g., [72, 75]). Additional genetic data (microsatellites, SNPs) can test whether they represent distinct evolutionary lineages.

### 4.5 Geographical distributions

The geographical ranges of the species vary considerably, and a diversity of processes seem to best explain their present distributions. Interestingly no speciation along a continental margin is observed in the complex.

The close phylogenetic relationship of `WA` with `EP` and Gala indicates that the closure of the isthmus of Panama led to the isolation of the species in the two basins.

The estimated ages of `Medit`, and the two Red Sea species are older (8.4-10.5 My, 1.1-1.7 My and 3.2-4.1 My respectively) than the last salinity crises characterizing these seas (*c*. 5 My for the Mediterranean [76] and potentially as early as *c*. 19,000 years ago for the Red Sea). This, in combination with the occurrence of these species outside their basins (`Medit` is known from Portugal and the Canaries; and both Red Sea species from Djibouti) indicate that these species invaded these seas recently and found refugia at the periphery of these basins during the salinity crises [77]. It is also interesting to note that the age of divergence for the two species from the Red Sea do not coincide suggesting that different events were responsible for the geographic isolation of these populations.

`tiger` and `tigerRedSea` are nocturnal species found on the reef slope. They are very sensitive to light, and retract quickly, deep into the reef matrix if a diver shines light on them. These species are known from a handful of individuals, and it is possible that their ranges are larger than what is reported here.

For species with wide ranges the causes of population divergence are less obvious, and the current distributions may not reflect species distributions at the time of speciation. The two most widespread species (`ESU1` and `ESU2`) are sympatric or parapatric with their closely related species, and both form the terminal clades in their respective radiations. This pattern suggests that these species were the sequential source of the peripheral species [7].

The lack of genetic differentiation between the Indian and Pacific Oceans for the species of the complex that occur on each sides of the Indo-Pacific barrier (IPB) is noteworthy. The IPB results from the restricted seaway between the Sunda and the Sahul shelves. During glaciation events, when sea levels were 100+ m below present level, only a narrow channel connected Indian and Pacific Oceans. In a recent literature survey, 15 out of 18 species of fish and invertebrate investigated showed genetic differentiation across the IBP associated with these low stands events [78]. Additionally, the genetic diversity of `ESU1` is low: among the 61 individuals sequenced, we found a total of 18 haplotypes, with the 3 most common haplotypes found in 40 individuals. Finally, both Marquesas and Oman, characterized by distinct ecological conditions from the rest of the Indo-Pacific hypothesized to increase the speed and probability of divergence, harbor the most widespread haplotype suggesting recent colonization of these peripheral areas to `ESU1`’s range. Together, these lines of evidence suggest a recent geographical expansion for the widespread species in the complex, but more analyses to confirm this hypothesis are needed.

Differences in the geographical ranges for some of the species might be explained by differences in habitat preferences. For instance, `tiger` is only found in the oligotrophic shallow waters of Micronesia and `Hawaii`, while `ESU3` and `gracilis` appear restricted to continental areas. For instance, `ESU3` found in Madagascar and Tanzania, was not encountered in the oceanic Mayotte or Scattered islands. We lack data to evaluate the role played by the ecology of the species in the diversification process and the co-existence of multiple species in sympatry.

### 4.6 Rates of secondary sympatry

Local diversity results from the accumulation of species that are reproductively isolated. Because reproductive isolation is most commonly initiated in allopatry [79], understanding how quickly species can be found in sympatry after diverging in geographical isolation is key to gain insights into the temporal dynamics of local diversity.

The restriction of marine invertebrates to “oceanic” and “continental” habitats has been documented for a range of organisms (e.g., land snails [80], marine snails [81, 82, 83], hermit crabs [7]), and a variety of biotic and abiotic factors may explain this pattern (reviewed in [7]). Interestingly, clades associated with continental habitats are characterized by much faster rates of secondary sympatry than species associated with insular habitats in both vertebrates (e.g., [84, 85]), and invertebrates (e.g., [86, 7, 83]). As a result, closely related species typically show strict allopatric distributions in oceanic setting (e.g., as in [86]) but often occur in sympatry along continental margins (e.g., [83]). Differences in rates of secondary sympatry vary across taxa but are faster in continental settings. For instance, in the wrasse *Anampes* sympatric sister species in continental setting diverged 600 ka ago, while the most closely related sympatric species in oceanic setting shared a common ancestor more than 10 Ma ago [85]. In the turbinid *Lunella*, none of the 7 lineages inhabiting oceanic islands that have diverged more than 15 Ma occur in sympatry while sister species found along the coastline of Oman have diverged 5 Ma [83].

The “*Holothuria impatiens*” complex shows a strickingly different pattern, and `ESU1, Hawaii` and `Wpac` are, to our knowledge, the fastest documented examples of speciation for a broadcast spawner invertebrate in an oceanic setting (`ESU1` and `Hawaii` diverged less than 2 Ma, and `ESU1` and `Wpac` about 2.5 Ma). Thus, species have been able to evolve reproductive isolation rapidly, and differ enough ecologically to allow for co-existence. In other words, the temporal dynamics of diversification in some oceanic species of the “*Holothuria impatiens*” complex is similar to what is observed for species occurring in continental settings.

Gamete recognition proteins (GRPs) play an important role in driving reproductive isolation in many free spawning organisms, such as sea urchins [87, 88] and gastropods [89]. For instance, strong positive selection has been detected in the GRPs between closely related species of sea urchins in sympatry but not in allopatry [90], suggesting the role of these proteins in maintaining species limits. GRPs have not yet been identified in sea cucumbers but bindin is known from sea urchins [88] and sea stars [91] which indicates that it is plesiomorphic in the clade that sea cucumbers emerged from. They could explain how species in the “*H. impatiens*” complex acquired reproductive isolation quickly.

Hellberg [92] proposed that climatic fluctuations isolate populations temporarily leading to speciation (transient allopatry), and as climatic conditions change, species shift their ranges. In continental settings, these shifts will lead to species coexistence as their distributions are constrained by the one dimensionality of the coastline. In insular conditions, however, these shifts in species distributions could be accompanied with the colonization of new islands and species could remain allopatric. Another difference is that in the continental setting the environmental conditions will vary continuously along the coast. Therefore, if species are adapted to different temperatures, the temperature gradient along the coast will allow species to overlap. However, in an insular setting, the absence of available habitat between the islands restricts the distribution of the species. These hypotheses could explain why most species remain allopatric for long periods of time and are restricted to archipelagos (as observed for `Hawaii` and `Gala`): rare migrants fail to colonize as local adaptation to environmental conditions render them maladapted to new islands. This model implies niche conservatism: species are narrowly adapted to a particular environment. The predictions of this model would suggest that `ESU1` has broadened its niche to allow the colonization of the range of `Hawaii` and `Wpac`.

### 4.7 Future directions

A shortcoming of the approach used in this study is that the results might be strongly influenced by the loci that harbor the most genetic differentiation, here the mitochondrial data. A recent extension of these methods, using single-nucleotide polymorphisms across the genome [93] would allow the confirmation of the species delineated in this study.

## 5 Conclusions

This study showed that the widespread, common and well known sea cucumber, *Holothuria impatiens* is actually a complex of at least 13 species. This estimate is conservative as additional lineages that could not be fully evaluated were also unraveled.

The multispecies coalescent framework allows the investigation of species limits in situations where evidence might have been inconclusive in the past. It is now possible to investigate species limits in groups that show some level of genetic differentiation but no morphological differences. The re-evaluation of species limits, and a better accuracy in the estimation of the divergence times, may improve our understanding of the speciation process by revealing new patterns and generating new hypotheses. Here, we showed that some species in this complex are characterized by the most rapid rate of secondary sympatry documented for a broadcast spawner in an oceanic setting. These results would need to be confirmed with genetic data that capture a larger fraction of the genome.

## References

[1] Pimm SL, Jenkins CN, Abell R, Brooks TM, Gittleman JL, et al. (2014) The biodiversity of species and their rates of extinction, distribution, and protection. Science (New York, NY) 344: 1246752.

[2] Appeltans W, Ahyong ST, Anderson G, Angel MV, Artois T, et al. (2012) The magnitude of global marine species diversity. Current biology 22: 2189–202.

[3] Costello MJ, May RM, Stork NE (2013) Can we name Earth’s species before they go extinct? Science (New York, NY) 339: 413–6.

[4] Moritz C (2002) Strategies to protect biological diversity and the evolutionary processes that sustain it. Systematic biology 51: 238–54.

[5] Knowlton N (1993) Sibling species in the sea. Annual Review of Ecology and Systematics 24: 189–216.

[6] Brown DM, Brenneman Ra, Koepfli KP, Pollinger JP, Milá B, et al. (2007) Extensive population genetic structure in the giraffe. BMC biology 5: 57.

[7] Malay MCMD, Paulay G (2010) Peripatric speciation drives diversification and distributional pattern of reef hermit crabs (Decapoda: Diogenidae: Calcinus). Evolution; international journal of organic evolution 64: 634–62.

[8] Prada C, Hellberg ME (2013) Long prereproductive selection and divergence by depth in a Caribbean candelabrum coral. Proceedings of the National Academy of Sciences of the United States of America 110: 3961–6.

[9] Hebert PDN, Penton EH, Burns JM, Janzen DH, Hallwachs W (2004) Ten species in one: DNA barcoding reveals cryptic species in the neotropical skipper butterfly Astraptes fulgerator. Proceedings of the National Academy of Sciences of the United States of America 101: 14812–7.

[10] Moritz C (1994) Defining ’Evolutionarily Significant Units’ for conservation. Trends in ecology & evolution 9: 373–5.

[11] Fujita MK, Leaché AD, Burbrink FT, McGuire Ja, Moritz C (2012) Coalescent-based species delimitation in an integrative taxonomy. Trends in ecology & evolution 27: 480–8.

[12] Baele G, Lemey P, Bedford T, Rambaut A, Suchard Ma, et al. (2012) Improving the accuracy of demographic and molecular clock model comparison while accommodating phylogenetic uncertainty. Molecular biology and evolution 29: 2157–67.

[13] Mayr E (1954) Geographic Speciation in Tropical Echinoids. Evolution 8: 1–18.

[14] Bowen BW, Rocha La, Toonen RJ, Karl Sa (2013) The origins of tropical marine biodiversity. Trends in Ecology & Evolution: 1–8.

[15] Forsska l P, Niebuhr C (1775) Descriptiones animalium, avium, amphibiorum, piscium, insectorum, vermium. ex officina Mölleri, 183 pp.

[16] Pearson J (1915) Notes on the Holothurioidea of the Indian Ocean. Spolia Zeylanica IX: 49–190.

[17] Rowe FWE (1969) A review of the family Holothuriidae (Holothurioidea: Aspidochirotida). Bulletin of the British Museum (Natural History) Zoology 18: 117–170.

[18] Harriott VJ (1985) Reproductive biology of three congeneric sea cucumber species, Holothuria atra, H. impatiens and H. edulis, at Heron Reef, Great Barrier Reef. Marine and Freshwater Research 36: 51.

[19] Flammang P, Ribesse J, Jangoux M (2002) Biomechanics of adhesion in sea cucumber Cuvierian tubules (Echinodermata, Holothuroidea)….and comparative biology 42: 1107–1115.

[20] Becker P, Flammang P (2010) Unravelling the sticky threads of sea cucumbers–A comparative study on Cuvierian tubule morphology and histochemistry. In: Byern J, Grunwald I, editors, Biological Adhesive Systems, Springer Vienna, chapter 6. pp. 87–98. doi:10.1007/978-3-7091-0286-2_6.

[21] Bakus G (1974) Toxicity in holothurians: a geographical pattern. Biotropica 6: 229–236.

[22] Roberts D, Bryce C (1982) Further observations on tentacular feeding mechanisms in holothurians. Journal of Experimental Marine Biology and Ecology 59: 151–163.

[23] Martens E, Clerck GD (1994) Interstitial and parasitic Platyhelminthes from the coast of the Seychelles. In: van der Land J, editor, Oceanic Reefs of the Seychelles, Leiden: National Museum of Natural History, chapter 6.5. pp. 1–10.

[24] Hampton JS (1958) Chemical Analysis of Holothurian Sclerites. Nature 181: 1608–1609.

[25] Hampton JJS (1959) Statistical analysis of holothurian sclerites. Micropaleontology 5: 335–349.

[26] Cutress B (1996) Changes in dermal ossicles during somatic growth in Caribbean littoral sea cucumbers (Echinoidea: Holothuroidea: Aspidochirotida). Bulletin of Marine Science 58: 44116.

[27] Lacey KMJ, McCormack GP, Keegan BF, Powell R (2005) Phylogenetic relationships within the class holothuroidea, inferred from 18S rRNA gene data. Marine Biology 147: 1149–1154.

[28] Honey-Escandon M, Laguarda-Figueras A, Solis-Marin FA (2012) Molecular phylogeny of the subgenus Holothuria (Selenkothuria) Deichmann, 1958 (Holothuroidea: Aspidochirotida). Zoological Journal of the Linnean Society 165: 109–120.

[29] Toral-Granda V (2008) Population status, fisheries and trade of sea cucumbers in Latin America and the Caribbean. In: Toral-Granda V, Lovatelli A, Vasconcelos MFDE, editors, Sea cucumbers. A global overview of fisheries and trade. FAO Fisheries and Aquaculture Technical Paper 516., Rome: FAO of the United Nations. pp. 213–229.

[30] Conand C, Muthiga NA (2007) Commercial sea cucumbers: a review for the Western Indian Ocean. WIOMSA Book Series 5: 66.

[31] Pakoa K, Lasi F, Tardy E, Friedman K (2009) The status of Sea Cucumbers exploited by Palau’s Subsistence Fishery. Nouméa, New Caledonia: Secretariat of the Pacific Community, 44 pp.

[32] Clark HL (1921) The Echinoderm fauna of the Torres Strait: its composition and its orgins, volume X of Department of Marine Biology of the Carnegie Institution of Washington. Washington D.C.: Carnegie Institution of Washington, 223 + 38 pl pp.

[33] Rowe F, Richmond M (2004) A preliminary account of the shallow-water echinoderms of Rodrigues, Mauritius, western Indian Ocean. Journal of Natural History 38: 3273–3314.

[34] Leaché AD, Fujita MK (2010) Bayesian species delimitation in West African forest geckos (Hemidactylus fasciatus). Proceedings Biological sciences / The Royal Society 277: 3071–7.

[35] Satler JD, Carstens BC, Hedin M (2013) Multilocus species delimitation in a complex of morphologically conserved trapdoor spiders (mygalomorphae, antrodiaetidae, aliatypus). Systematic biology 62: 805–23.

[36] Grummer Ja, Bryson RW, Reeder TW (2014) Species Delimitation Using Bayes Factors: Simulations and Application to the Sceloporus scalaris Species Group (Squamata: Phrynosomatidae). Systematic biology 63: 119–33.

[37] Pons J, Barraclough T, Gomez-Zurita J, Cardoso A, Duran D, et al. (2006) Sequence-Based Species Delimitation for the DNA Taxonomy of Undescribed Insects. Systematic Biology 55: 595–609.

[38] Monaghan MT, Wild R, Elliot M, Fujisawa T, Balke M, et al. (2009) Accelerated species inventory on Madagascar using coalescent-based models of species delineation. Systematic biology 58: 298–311.

[39] Heled J, Drummond A (2010) Bayesian inference of species trees from multilocus data. Molecular biology and evolution 27: 570–580.

[40] Sonnenberg R, Nolte AW, Tautz D (2007) An evaluation of LSU rDNA D1-D2 sequences for their use in species identification. Frontiers in zoology 4: 6.

[41] Biomatters Ltd (2012). Geneious 5.5.8.

[42] Hoareau TB, Boissin E (2010) Design of phylum-specific hybrid primers for DNA barcoding: addressing the need for efficient COI amplification in the Echinodermata. Molecular ecology resources 10: 960–7.

[43] Arndt A, Marquez C, Lambert P, Smith MJ (1996) Molecular phylogeny of eastern Pacific sea cucumbers (Echinodermata: Holothuroidea) based on mitochondrial DNA sequence. Molecular phylogenetics and evolution 6: 425–37.

[44] Fujisawa T, Barraclough TG (2013) Delimiting species using single-locus data and the generalized mixed yule coalescent approach: a revised method and evaluation on simulated data sets. Systematic biology 62: 707–24.

[45] Drummond AJ, Bouckaert RR (2014) Bayesian evolutionary analysis with BEAST 2. Cambridge University Press.

[46] Talavera G, Dincă V, Vila R (2013) Factors affecting species delimitations with the GMYC model: insights from a butterfly survey. Methods in Ecology and Evolution 4: 1101–1110.

[47] Kingman J (1982) The coalescent. Stochastic Processes and their Applications 13: 235–248.

[48] Griffiths RC, Tavaré S (1994) Sampling theory for neutral alleles in a varying environment. Philosophical transactions of the Royal Society of London Series B, Biological sciences 344: 40310.

[49] Yule G (1925) A mathematical theory of evolution, based on the conclusions of Dr. JC Willis, FRS. Philosophical Transactions of the Royal Society of London Series B, Containing Papers of a Biological Character 213: 21–87.

[50] Drummond AJ, Suchard Ma, Xie D, Rambaut A (2012) Bayesian phylogenetics with BEAUti and the BEAST 1.7. Molecular biology and evolution 29: 1969–73.

[51] Ayres DL, Darling A, Zwickl DJ, Beerli P, Holder MT, et al. (2012) BEAGLE: an application programming interface and high-performance computing library for statistical phylogenetics. Systematic biology 61: 170–3.

[52] Lanfear R, Calcott B, Ho SYW, Guindon S (2012) Partitionfinder: combined selection of partitioning schemes and substitution models for phylogenetic analyses. Molecular biology and evolution 29: 1695–701.

[53] Lessios H (2008) The Great American Schism: Divergence of Marine Organisms After the Rise of the Central American Isthmus. Annual Review of Ecology, Evolution, and Systematics 39: 63–91.

[54] Heled J, Bouckaert RR (2013) Looking for trees in the forest: summary tree from posterior samples. BMC evolutionary biology 13: 221.

[55] Ezard T, Fujisawa T, Barraclough T (2013) splits: SPecies’ LImits by Threshold Statistics (R package version 1.0-19/r48) http://r-forge.r-project.org/projects/splits/.

[56] Kass RE, Raftery AE (1995) Bayes Factors. Journal of the American Statistical Association 90: 773–795.

[57] Edgar RC (2004) MUSCLE: multiple sequence alignment with high accuracy and high throughput. Nucleic acids research 32: 1792–7.

[58] Castresana J (2000) Selection of conserved blocks from multiple alignments for their use in phylogenetic analysis. Molecular biology and evolution 17: 540–52.

[59] Stamatakis A (2014) RAxML version 8: a tool for phylogenetic analysis and post-analysis of large phylogenies. Bioinformatics (Oxford, England) 30: 1312–3.

[60] Bouckaert R, Heled J, Kühnert D, Vaughan T, Wu CH, et al. (2014) BEAST 2: A Software Platform for Bayesian Evolutionary Analysis. PLoS computational biology 10: e1003537.

[61] Hasegawa M, Kishino H, Yano T (1985) Dating of the human-ape splitting by a molecular clock of mitochondrial DNA. Journal of molecular evolution 22: 160–74.

[62] Papadopoulou A, Bergsten J, Fujisawa T, Monaghan MT, Barraclough TG, et al. (2008) Speciation and DNA barcodes: testing the effects of dispersal on the formation of discrete sequence clusters. Philosophical transactions of the Royal Society of London Series B, Biological sciences 363: 2987–96.

[63] Esselstyn Ja, Evans BJ, Sedlock JL, Anwarali Khan FA, Heaney LR (2012) Single-locus species delimitation: a test ofthe mixed Yule-coalescent model, with an empirical application to Philippine round-leaf bats. Proceedings Biological sciences / The Royal Society 279: 3678–86.

[64] Ratnasingham S, Hebert PDN (2013) A DNA-based registry for all animal species: the barcode index number (BIN) system. PloS one 8: e66213.

[65] Puillandre N, Lambert a, Brouillet S, Achaz G (2012) ABGD, Automatic Barcode Gap Discovery for primary species delimitation. Molecular ecology 21: 1864–77.

[66] Zhang J, Kapli P, Pavlidis P, Stamatakis A (2013) A general species delimitation method with applications to phylogenetic placements. Bioinformatics (Oxford, England) 29: 2869–76.

[67] Reid NM, Carstens BC (2012) Phylogenetic estimation error can decrease the accuracy of species delimitation: a Bayesian implementation of the general mixed Yule-coalescent model. BMC evolutionary biology 12: 196.

[68] Marshall DC, Hill KBR, Cooley JR, Simon C (2011) Hybridization, mitochondrial DNA phylogeography, and prediction of the early stages of reproductive isolation: lessons from New Zealand cicadas (genus Kikihia). Systematic biology 60: 482–502.

[69] Leache AD, Harris RB, Rannala B, Yang Z (2013) The Influence of Gene Flow on Species Tree Estimation: A Simulation Study. Systematic biology 0: 1–14.

[70] Harris RB, Carling MD, Lovette IJ (2013) the Influence of Sampling Design on Species Tree Inference: a New Relationship for the New World Chickadees (Aves: Poecile). Evolution; international journal of organic evolution.

[71] Clark HL (1938) Memoirs of Comparative Zoology at Harvard College. Vol LV. Echinoderms from Australia an account of collections made in 1929 and 1938. Cambridge, MA, USA: Museum of Comparative Zoology, 596 + 28 plates pp.

[72] McMillan WO, Weigt LA, Palumbi SR (1999) Color Pattern Evolution, Assortative Mating, and Genetic Differentiation in Brightly Colored Butterflyfishes (Chaetodontidae). Evolution 53: 247.

[73] Detto T (2007) The fiddler crab Uca mjoebergi uses colour vision in mate choice. Proceedings Biological sciences / The Royal Society 274: 2785–90.

[74] Baldwin J, Johnsen S (2012) The male blue crab, Callinectes sapidus, uses both chromatic and achromatic cues during mate choice. The Journal of experimental biology 215: 1184–91.

[75] Gaither MR, Schultz JK, Bellwood DR, Pyle RL, Dibattista JD, et al. (2014) Evolution of pygmy angelfishes: recent divergences, introgression, and the usefulness of color in taxonomy. Molecular phylogenetics and evolution 74: 38–47.

[76] Hernandez-Molina FJ, Stow DaV, Alvarez-Zarikian Ca, Acton G, Bahr a, et al. (2014) Onset of Mediterranean outflow into the North Atlantic. Science 344: 1244–1250.

[77] DiBattista JD, Berumen ML, Gaither MR, Rocha La, Eble Ja, et al. (2013) After continents divide: comparative phylogeography of reef fishes from the Red Sea and Indian Ocean. Journal of Biogeography 40: 1170–1181.

[78] Gaither MR, Toonen RJ, Robertson DR, Planes S, Bowen BW (2010) Genetic evaluation of marine biogeographical barriers: perspectives from two widespread Indo-Pacific snappers (Lutjanus kasmira and Lutjanus fulvus). Journal of Biogeography 37: 133–147.

[79] Coyne JA, Orr HA (2004) Speciation. Sunderland, MA, USA: Sinauer Associates, Inc.

[80] Paulay G (1994) Biodiversity on oceanic islands: its origin and extinction. American Zoologist 34: 134–144.

[81] Abbott RT (1960) The genus Strombus in the Indo-Pacific. Indo-Pacific Mollusca Monographs of the marine mollusks of the tropical West Pacific and Indian Oceans 1: 33–146.

[82] Reid DG, Lal K, Mackenzie-Dodds J, Kaligis F, Littlewood DTJ, et al. (2006) Comparative phylogeography and species boundaries in Echinolittorina snails in the central Indo-West Pacific. Journal of Biogeography 33: 990–1006.

[83] Williams ST, Apte D, Ozawa T, Kaligis F, Nakano T, et al. (2011) Speciation and dispersal along continental coastlines and island arcs in the Indo-West Pacific turbinid gastropod genus Lunella. Evolution; international journal of organic evolution 65: 1752–71.

[84] Taylor MS, Hellberg ME (2005) Marine Radiations at Small Geographic Scales: Speciation in Neotropical Reef Gobies (Elacatinus). Evolution 59: 374–385.

[85] Hodge JR, Read CI, van Herwerden L, Bellwood DR (2012) The role of peripheral endemism in species diversification: evidence from the coral reef fish genus Anampses (Family: Labridae). Molecular phylogenetics and evolution 62: 653–63.

[86] Meyer CP, Geller JB, Paulay G (2005) Fine scale endemism on coral reefs: archipelagic differentiation in turbinid gastropods. Evolution; international journal of organic evolution 59: 113–25.

[87] Levitan DR, Ferrell DL (2006) Selection on gamete recognition proteins depends on sex, density, and genotype frequency. Science (New York, NY) 312: 267–9.

[88] Lessios HA (2011) Speciation Genes in Free-Spawning Marine Invertebrates. Integrative and comparative biology 51: 456–465.

[89] Hellberg ME, Moy GW, Vacquier VD (2000) Positive selection and propeptide repeats promote rapid interspecific divergence of a gastropod sperm protein. Molecular biology and evolution 17: 458–66.

[90] Geyer LB, Palumbi SR (2003) Reproductive character displacement and the genetics of gamete recognition in tropical sea urchins. Evolution; international journal of organic evolution 57: 1049–60.

[91] Patiño S, Aagaard JE, MacCoss MJ, Swanson WJ, Hart MW (2009) Bindin from a sea star. Evolution & development 11: 376–81.

[92] Hellberg ME (1998) Sympatric sea shells along the sea’s shore: the geography of speciation in the marine gastropod Tegula. Evolution 52: 1311–1324.

[93] Leaché AD, Fujita MK, Minin VN, Bouckaert RR (2014) Species Delimitation using Genome-Wide SNP Data. Systematic biology 63: 534–42.

